# Conserved cysteines in titin sustain the mechanical function of cardiomyocytes

**DOI:** 10.1101/2020.09.05.282913

**Authors:** Elías Herrero-Galán, Fernando Domínguez, Inés Martínez-Martín, Cristina Sánchez-González, Natalia Vicente, Laura Lalaguna, Elena Bonzón-Kulichenko, Enrique Calvo, Esther González-López, Marta Cobo-Marcos, Belén Bornstein, Ana Briceño, Juan Pablo Ochoa, Jose Maria Garcia-Aznar, Carmen Suay-Corredera, Maria Rosaria Pricolo, Ángel Fernández-Trasancos, Diana Velázquez-Carreras, Claudio Badía Careaga, Belén Prados, Francisco Gutiérrez-Agüera, Mahmoud Abdellatif, Simon Sedej, Peter P. Rainer, David Giganti, Giovanna Giovinazzo, Juan A. Bernal, Raúl Pérez-Jiménez, Torsten Bloch Rasmussen, Thomas Morris Hey, Inmaculada Vivo-Ortega, Jesús Piqueras-Flores, Enrique Lara-Pezzi, Jesús Vázquez, Pablo Garcia-Pavia, Jorge Alegre-Cebollada

## Abstract

The protein titin determines cardiomyocyte contraction and truncating variants in the titin gene (*TTN*) are the most common cause of dilated cardiomyopathy (DCM). Different to truncations, missense variants in *TTN* are currently classified as variants of uncertain significance due to their high frequency in the population and the absence of functional annotation. Here, we report the regulatory role of conserved, mechanically active titin cysteines, which, contrary to current views, we uncover to be reversibly oxidized in basal conditions leading to isoform- and force-dependent modulation of titin stiffness and dynamics. Building on our functional studies, we demonstrate that missense mutations targeting a conserved titin cysteine alter myocyte contractile function and cause DCM in humans. Our findings have a direct impact on genetic counselling in clinical practice.

**One sentence summary:** Mutations targeting cysteines key to the mechanoredox control of titin cause human dilated cardiomyopathy

## INTRODUCTION

Titin is a fundamental protein for the contractile function of striated myocytes since it provides structural support and sets the stiffness of sarcomeres (*1–3*). Not surprisingly, the mechanical properties of titin are exquisitely modulated both transcriptionally through specific alternative mRNA splicing (*4, 5*), and posttranslationally via biochemical modifications such as phosphorylation (*6*). Truncating variants in the titin gene (*TTN*) are the main cause of dilated cardiomyopathy (DCM), a disease that is the most frequent trigger of heart failure in the young and of heart transplantation worldwide (*7–11*). However, the primordial pathomechanisms remain elusive, reflecting our incomplete knowledge of the function of titin (*12–14*). Akin to other sarcomeric proteins in which both truncations and missense variants lead to cardiomyopathy (*9*), it has been speculated that rare missense mutations in *TTN* could be the cause of DCM in some of the ~50% genotype-negative patients (*7, 15, 16*). Indeed, numerous titin missense variants can be identified in DCM individuals, although at a frequency that is not significantly higher than in the general population (*17*). Hence, when rare missense variants are identified in *TTN*, they are currently classified as variants of unknown significance. It is conceivable though that population genetics analyses have limited power to set apart true pathogenic mutations from a background of phenotypically silent *TTN* missense variants. Here, we propose an alternative approach by which we first identify conserved residues that are important for the function of titin, which we then screen for pathogenic missense variants in DCM patients. We have focused on a group of structurally conserved cysteine residues whose modification has been suggested, on the basis of *in vitro* evidence, to modulate the mechanical properties of titin (*18*). Following our strategy, we show the *in vivo* relevance of cysteine-based regulation of titin mechanics and unequivocally demonstrate that rare *TTN* missense variants cause DCM in humans.

## RESULTS

### Titin cysteines are reversibly oxidized in basal conditions

The mechanical properties of titin stem from force-dependent conformational changes of polypeptide regions that belong to the extensible I-band region of the protein (**Figure 1A**). These conformational changes include extension and entropic recoil of the serially linked immunoglobulin-like (Ig) domains and the random-coil N2Bus and PEVK regions, and Ig domain unfolding and refolding transitions (*19*). Interestingly, the I-band of titin is rich in cysteine residues (*18, 20*), many of which appear at structurally and evolutionary conserved positions within Ig domains (**Figure 1B-F, Supplementary Figure S1, Supplementary File S1**). *In vitro* experiments have shown that oxidation of these cysteines has major mechanical consequences. For instance, disulfide bonds established by the triad of conserved cysteines B, F and G in Ig domains (98% mean evolutionary conservation, **Figure 1B,C,E**) stiffen titin via reduction of the protein contour length (*21*), while S-thiolation of cysteines 47 and 63 (94% mean evolutionary conservation, **Figure 1B,D,E**) lead to titin softening through Ig folding inhibition (*22*). These five structurally conserved cysteines are more evolutionary conserved than cysteines appearing in other positions in Ig domains (97% *vs* 89% mean evolutionary conservation, **Figure 1F**). The N2Bus region also contains cysteines that can stiffen titin through disulfide bond formation (*23*), albeit they are less evolutionary conserved than Ig domain cysteines (53% mean conservation, **Figure 1F, Supplementary Figure S1C**). Beyond the remarkable conservation of I-band titin’s cysteine residues and their role in the evolution of the protein in vertebrates (*24*), the *in vivo* relevance of redox mechanical modulation of titin is supported by limited data on the global oxidation of the protein (*25–28*), the effects of redox-active molecules on striated muscle mechanics (*22, 23, 29, 30*), and the disulfide-compatible location of the majority of structurally conserved cysteines of titin (*21, 31*). However, the extent and location of native titin oxidations remain unexplored, limiting our understanding of the impact of oxidative modifications on the mechanical function of the protein.

**Figure 1.**
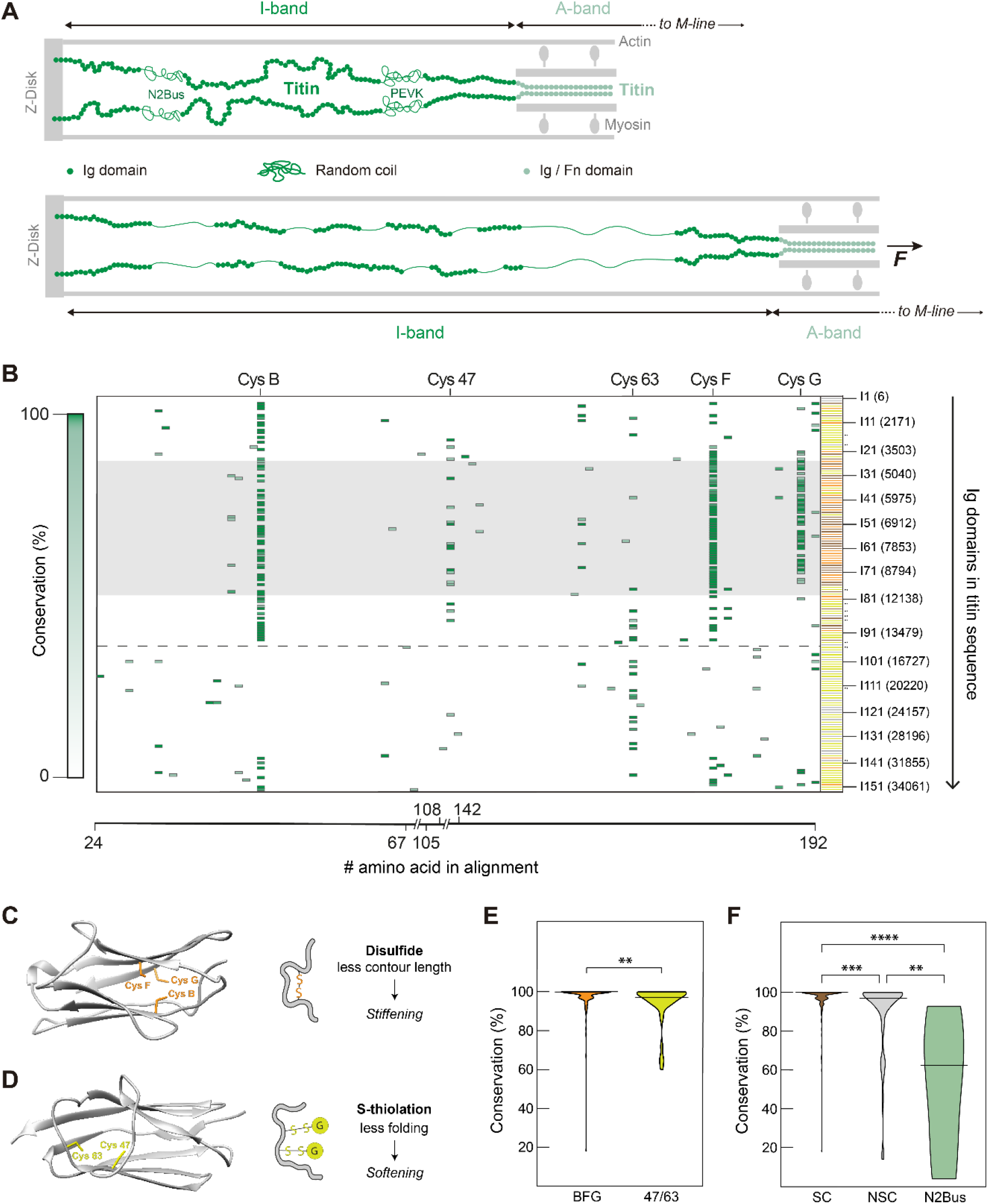
Evolutionary and structurally conserved cysteines in titin. **A**: *Top:* Representation of half a sarcomere (not to scale), indicating the positions of the I-band (green) and the A-band (teal) of titin. Immunoglobulin-like (Ig), random-coil (PEVK and N2Bus) and fibronectin III (Fn) domains are shown. *Bottom*: Under mechanical force, the serially linked Ig domains and the random coil regions extend, and Ig domains unfold. **B**: Map of cysteine positions along the alignment of Ig domains of human titin (breaks leave out Cys-free segments of the alignment). Structurally conserved positions B, 47, 63, F and G are indicated on top of the alignment. Positions are colored according to percentage of evolutionary conservation, from 0 (white) to 100% (dark green). The number and positions of Ig domains according to Uniprot Q8WZ42-1 (N2BA isoform) are indicated on the right. Pairs of dots mark Ig domains that are not annotated in Uniprot (*21*). Ig domains are represented in grey if they are cysteine-free, in orange if they contain at least two of the triad, disulfide-competent cysteines B, F and G (*21*), in yellow if they contain cysteines but no more than 1 cysteine B, F or G, and in brown if at least two triad cysteines are present together with other cysteines. Spliced-out region in the N2B titin isoform is indicated by the shaded grey area. The horizontal dashed line shows the boundary between the I- and A-bands of titin (positions 14018-14019). **C**: 3D homology model of an Ig domain containing the disulfide-competent CysB-CysF-CysG triad (I74, positions 9079-9168 from Uniprot entry Q8WZ42). The reduced contour length of the disulfide-containing unfolded state leads to domain stiffening. **D**: High-resolution structure of the traditionally named I27 domain of titin (also known as I91, PDB code 1tit), which contains unpaired, S-thiolation-competent cysteines 47 and 63 (positions 12674-12765 from Uniprot entry Q8WZ42). S-thiolations inhibit folding, leading to domain softening (*22*). **E**: Violin-plot distribution of percentages of evolutionary conservation of structurally conserved BFG (n=153) and 47/63 (n=40) cysteines in human titin. p(BFG *vs* 47/63)=0.0063 (Mann-Whitney). **F**: Violin-plot distribution of percentages of evolutionary conservation of structurally conserved (SC, including BFG and 47/63 cysteines, n=193), non-structurally conserved (NSC, n=113), and N2Bus (n=6) cysteines in human titin. p(SC *vs* NSC)=0.0001, p(SC *vs* N2Bus)<0.0001, p(NSC *vs* N2Bus)=0.0033 (Kruskal-Wallis and Dunn’s multiple comparisons test). Evolutionary conservation values in panels E and F were calculated from the alignment of 36 titin sequences from different species (**Supplementary Figure S1 and Supplementary File S1**). Horizontal bars in violin plots indicate median values.

To measure the oxidation state of native titin, we first developed an SDS-PAGE-based assay that exploits thiol chemistry reactions to block reduced cysteines with an alkylating agent and label reversibly oxidized cysteines with the fluorophore monobromobimane (mBBr) (**Figure 2A**) (*32*). The assay takes advantage of unequivocal identification of titin as the slowest-migrating protein in low-percentage-acrylamide SDS-PAGE gels (*33, 34*). We verified that the mBBr fluorescence signal of titin cysteines is linearly dependent on the amount of lysate analyzed (**Figure 2B, Supplementary Figure S2**) and used Coomassie staining (**Figure 2C, Supplementary Figure S2**) to get normalized oxidation measurements that are independent of the amount of protein loaded on the gel (**Figure 2D**). The initial alkylating step is done in denaturing conditions to block all reduced cysteines. To limit artifactual oxidation signal, this step needs to be fast and complete. We chose N-ethylmaleimide (NEM) as the alkylating agent because of its superior reaction kinetics and its high solubility in aqueous buffers (*27, 35*). Using murine cardiac samples, we found that the oxidation signal of titin plateaus when the concentration of NEM in the lysis buffer is above 5 mM (**Figure 2E**), suggesting that these conditions result in minimal artifactual oxidation. To further ensure efficient thiol blockage, in all our subsequent experiments we perfused myocardial tissue with PBS containing 50 mM NEM immediately after sacrifice. The same high NEM concentration was kept during lysis. As an additional preventive measure, we subtracted the mBBr signal of samples not incubated with DTT to account for potential reduced thiols refractory to NEM blockage (see Methods). Using our in-gel mBBr fluorescence assay, we observed that the extent of reversible cysteine oxidation of titin is ~4 times higher than that of myosin, a partner protein of titin in the sarcomere (**Figure 2F**), confirming that a fraction of titin cysteines are constitutively oxidized in cardiac tissue.

**Figure 2.**
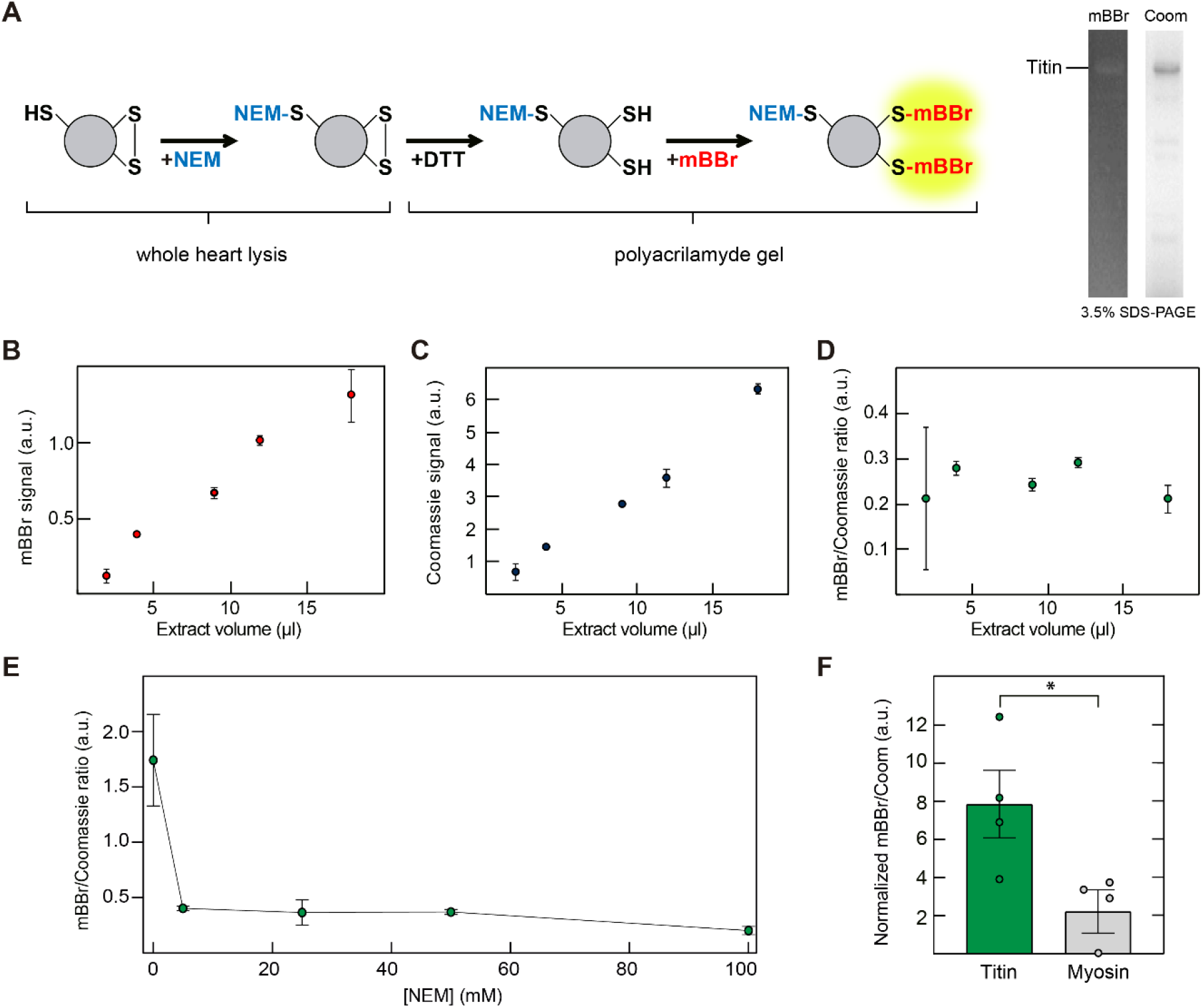
Titin is oxidized in basal conditions. **A**: *Left*: Reaction scheme to label reversibly oxidized thiols with mBBr. Protein extracts in which reduced thiols have been initially blocked by NEM are run in an SDS-PAGE gel. Following electrophoresis, oxidized thiols are reduced by DTT and then labelled with the fluorescent probe mBBr. *Right*: example SDS-PAGE gel used to quantify mBBr fluorescence from titin bands. Coomassie staining of the same gel is used for normalization. **B, C**: The mBBr and Coomassie signals originating from titin are linearly dependent on the amount of lysate loaded in SDS-PAGE gels (all data points are the average of duplicates) (**Supplementary Figure S2**). **D**: Final mBBr/Coomassie ratio is independent of the amount of lysate loaded in the gel. **E**: Scan to determine the concentration of NEM needed to completely block initially reduced titin thiols. Duplicates were performed for each NEM concentration. **F**: Quantification of the oxidation of titin and myosin in four different mouse hearts using 3.5% and 12% SDS-PAGE gels, respectively. The volume of lysate analyzed was adjusted so that both titin (15 μl) and myosin (3 μl) were in the linear range of detection for Coomassie. Oxidation signal is normalized by the density of cysteines of each protein (13.3 Cys/1000 amino acid for titin, 7.2 Cys/1000 amino acid for myosin, see Methods). n =4 animals, p=0.0286 (Mann-Whitney).

### Titin oxidation is boosted upon birth

Having demonstrated that titin is oxidized *in vivo*, we set out to test if titin oxidation is modulated physiologically. With this aim, we turned to the perinatal model. In mammals, birth is accompanied by exposure to high O_2_ concentration, which impacts on the function of many cells and organs including the heart (*36*). Indeed, during early postnatal development, cardiomyocytes arrest cell cycle following activation of oxidative signals (*37*). We hypothesized that increased oxidative conditions could result in higher levels of titin oxidation upon birth, which we set out to test using the in-gel mBBr fluorescence assay. In order to increase statistical power, in the experiments we included control proteins that enabled averaging data from multiple samples (**Supplementary Figure S3A**, see Methods). Since the proportion of titin isoforms in the heart changes during postnatal development (*4, 38*), and these isoforms have different proportion of cysteines (*21*), we normalized the results according to the cysteine density at each developmental stage (**Figure 3, Supplementary Figure S3B**, see Methods). Supporting our initial hypothesis, we found that cardiac titin oxidation is 43±15% (mean ± SEM) higher in newborn (P0) mice than in E18.5 embryos, while we detected no significant difference between P0 and adult samples.

**Figure 3.**
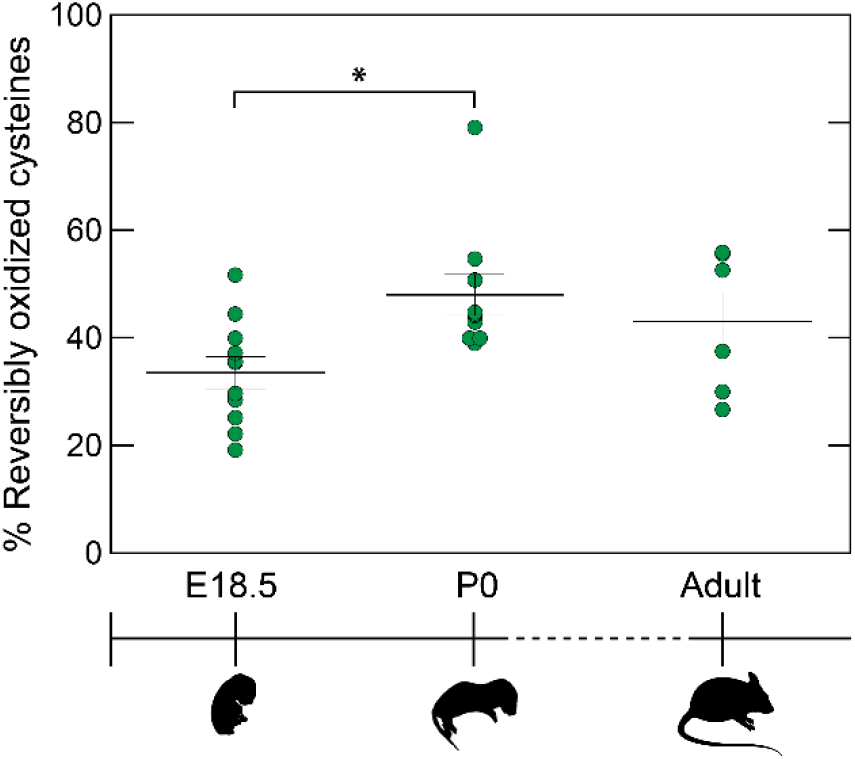
Titin oxidation is increased upon birth. Reversible oxidation of titin cysteines in samples from 18.5-days embryos (E18.5, n = 11), newborn (P0, n = 10) and adult mice (n = 6). Density of reversibly oxidized cysteines (**Supplementary Figure S3A**) is compared to the total density of cysteines according to the percentages of each titin isoform (14.6 Cys/1000 amino acid for N2BA and 13.3 Cys/1000 amino acid for N2B, **Supplementary Figure S3Bn**; see Methods). p(E18.5 *vs* P0) = 0.0259, p(E18.5 *vs* Adult) = 0.259, p(P0 *vs* Adult) > 0.999 (Kruskal-Wallis and Dunn’s multiple comparisons test).

### Preferential oxidation of I-band cysteines

To characterize the landscape of reversible titin cysteine oxidations, we resourced to mass spectrometry (MS). We analyzed P0 samples because they contain a high proportion of the cysteine-rich N2BA titin isoform. Fluorescent titin bands were sliced from SDS-PAGE gels and treated with trypsin. The resulting peptides were subjected to LC-MS analysis and MS/MS spectra were searched against a mouse proteome database. As expected from the full titin band separation in 3.5% SDS-PAGE gels, 79 ± 1% (3 replicates) of the identified species corresponded to titin-derived peptides, even when using a relaxed identification criterion (5% False Discovery Rate, FDR) (**Figure 4A**). Titin coverage in these searches was 33±3% (**Figure 4B**). To maximize the coverage of cysteine peptides, we followed an alternative, more targeted strategy (*39*). Since most detected peptides in the search against the full proteome originate from titin, we repeated the search against a database containing only titin and all identified cysteine peptides were validated using Vseq (*40*). Similar to the in-gel fluorescence method, we excluded mBBr-derivatized peptides when also detected in –DTT control samples, resulting in a final 45% aggregated cysteine coverage at a FDR<2% (see Methods).

**Figure 4.**
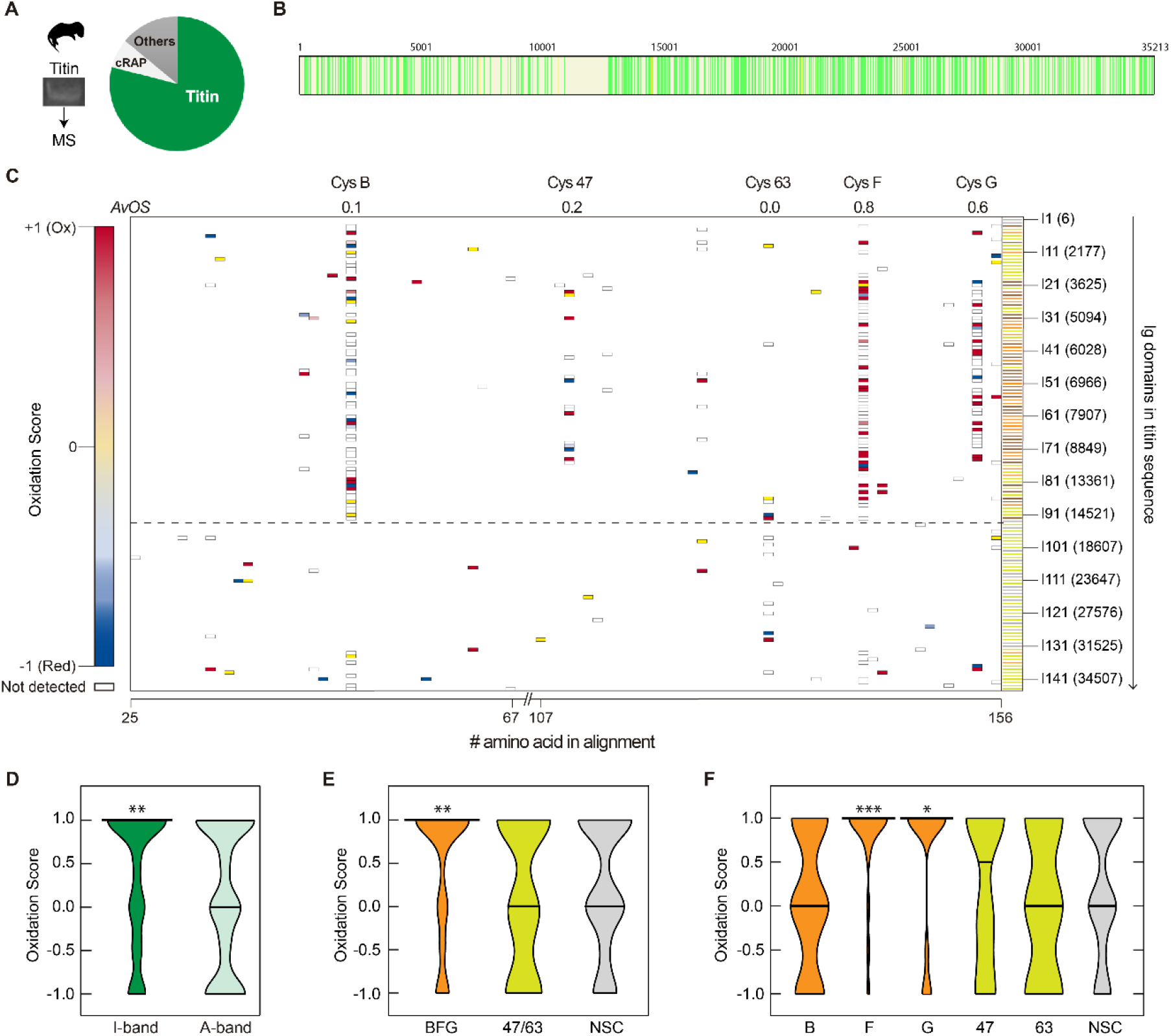
Landscape of cysteine oxidation in murine postnatal (P0) cardiac titin. **A**: Fraction of identified peptides belonging to titin (reference sequence Uniprot A2ASS6-1, which codes for the long N2BA titin isoform) when the MS/MS spectra are searched against a mouse proteome database containing common contaminants (cRAP). **B**: Aggregated itin coverage obtained after searching against the mouse proteome (n =3 samples; yellow and green, peptides detected with FDR<5% and <1%, respectively). **C**: Oxidation Score (OS) map for every cysteine position in the alignment of Ig domains of mouse titin, according to the color code indicated on the left (red, oxidized; blue, reduced). Average OS (AvOS) for cysteines 47, 63 and the BFG triad across all Ig domains are indicated. Cysteines shown in white were not identified and do not contribute to calculation of OS. The horizontal dashed line shows the boundary between the I- and A-bands of titin (positions 14880-14881 in Uniprot A2ASS6-1). The number and positions of Ig domains are indicated on the right. Ig domains are classified and colored as in Figure 1. **D**: Violin-plot distributions of OS for cysteines belonging to the I-(n=97) and A-(n=123) bands of murine titin. p(OS>0, I-band) = 0.0054, p(OS>0, A-band) = 0.9886 (hypergeometric test). **E**: Violin-plot distributions of OS for cysteines BFG (n=61), 47/63 (n=14) and non-structurally conserved (NSC, n=33). p(OS>0, BFG) = 0.0054, p(OS>0, 47/63) = 0.8137, p(OS>0, NSC) = 0.9268 (hypergeometric test). **F**: Violin-plot distributions of OS for cysteines B (n=21), F (n=23), G (n=17), 47 (n=8), 63 (n=6) and NSC (n=33). p(OS>0, B) = 0.9593, p(OS>0, F) = 0.0001, p(OS>0, G) = 0.0200, p(OS>0, 47) = 0.53365, p(OS>0, 63) = 0.7894, p(OS>0, NSC) = 0.9268 (hypergeometric test). Horizontal bars in violin plots indicate the median values.

In the MS experiments, we detect reduced and oxidized cysteines from the mass shifts associated to NEM and mBBr modifications, respectively (**Figure 2A**). To obtain a global picture of oxidation, we have calculated an oxidation score (OS) for each cysteine position in the alignment of the Ig domains of titin (**Figure 4C**, see Methods). OS data show that titin cysteines of the mechanically active I-band are significantly more oxidized than those belonging to the A-band (**Figure 4D**). In addition, cysteines of the disulfide-competent triad BFG (in particular cysteines F and G) are detected as oxidized more frequently than other cysteines in titin (**Figure 4C,E,F**). Unfortunately, we did not detect cysteine-containing peptides from the N2Bus region of murine titin, so its native oxidation state could not be determined.

### Titin oxidation in human hearts

We next examined the landscape of cysteine oxidation in human cardiac titin using left ventricular snap-frozen samples from two non-failing donor hearts. These samples were processed and analyzed by LC-MS following the same approach used for mouse samples. 74±3% of identified peptides were derived from titin when the MS/MS search was done against the human proteome database at a 5% FDR (**Figure 5A**). In these searches, titin coverage was 37±6% (**Figure 5B**). As in experiments using mouse samples, we run targeted searches against a database containing only human titin and the resulting cysteine peptides were validated with Vseq. mBBr-derivatized peptides also detected in – DTT samples were excluded from analysis, leading to a final 39% aggregated cysteine coverage at a FDR<2%. Results show similar levels of global oxidation in humans and in mice (global OS are 0.1 and 0.4, respectively, **Figure 5C**). As in mice, cysteines of the I-band are significantly more oxidized than those of the A-band (**Figure 5D**) and the OS of structurally conserved cysteines in Ig domains tends to be higher than that of non-structurally conserved cysteines (**Figure 5C,E**). Similar to results with murine samples, cysteine F shows the highest OS among disulfide-competent positions, although in human samples the I-band-specific cysteine 47 has the highest OS among the structurally conserved cysteines (**Figure 5C,F**). In one of the human samples, we detected two cysteines belonging to the N2Bus region (Cys4083 was detected as reduced, and Cys4124 was found in oxidized and reduced states). In summary, MS results indicate that reversible oxidations are also present in native titin from non-failing human myocardium.

**Figure 5.**
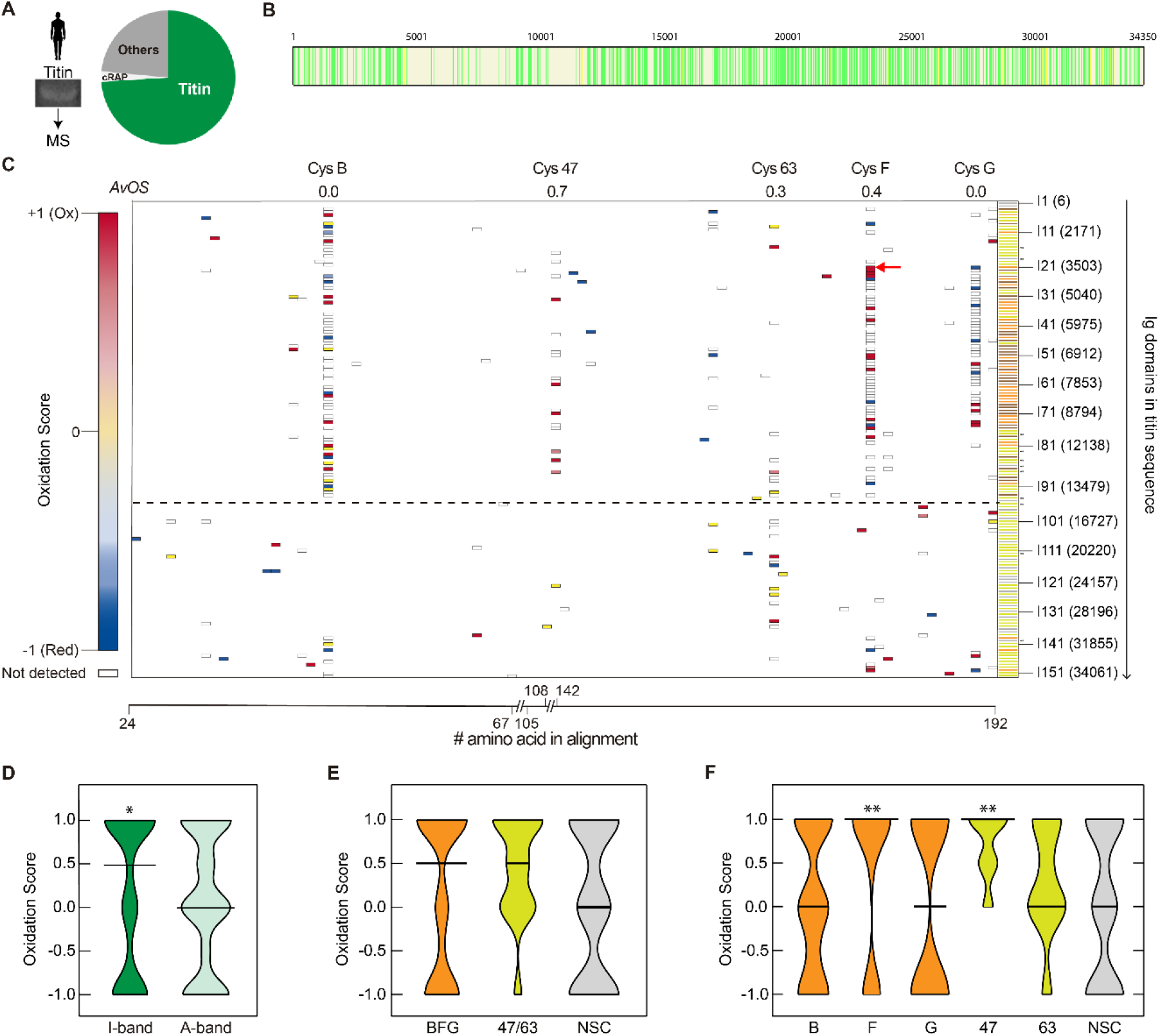
Landscape of cysteine oxidation in adult human cardiac titin. **A**: Fraction of identified peptides belonging to titin (reference sequence Uniprot Q8WZ42-1, which corresponds to the long N2BA titin isoform) when the MS/MS spectra are searched against a database containing the human proteome and common contaminants (cRAP). **B**: Aggregated titin coverage obtained after searching against the human proteome (n = 2 samples; yellow and green, peptides detected with FDR<5% and <1%, respectively). **C**: Oxidation Score (OS) map for every cysteine position in the alignment of Ig domains of human titin, according to the color code indicated on the left (red, oxidized; blue, reduced). Average OS (*AvOS*) for cysteines 47, 63 and the B/F/G triad across all Ig domains are indicated. Cysteines shown in white were not identified and do not contribute to calculation of OS. Red arrow points to Cys3575, a target of DCM-linked mutations. The dashed line shows the boundary between the I- and A-bands of titin (positions 14018-14019 in Uniprot Q8WZ42-1). The number and positions of Ig domains are indicated on the right. Pairs of dots mark Ig domains that are not annotated in Uniprot. Ig domains are classified and colored as in Figure 1. **D**: Violin-plot distributions of OS for cysteines belonging to the I-(n=77) and A-(n=122) bands of human titin. p(OS>0, I-band) = 0.0435, p(OS>0, A-band) = 0.9220 (hypergeometric test). **E**: Violin-plot distributions of oxidation scores for cysteines BFG (n=57), 47/63 (n=16) and non-structurally conserved (NSC, n=33). p(OS>0, BFG) = 0.2746, p(OS>0, 47/63) = 0.0745, p(OS>0, NSC) = 0.8705 (hypergeometric test). **F**: Violin-plot distributions of oxidation scores for cysteines B (n=25), F (n=20), G (n=12), 47 (n=7), 63 (n=9) and NSC (n=33) in human cardiac titin. p(OS>0, B) = 0.8974, p(OS>0, F) = 0.0093, p(OS>0, G) = 0.3537, p(OS>0, 47) = 0.0055, p(OS>0, 63) = 0.4759, p(OS>0, NSC) = 0.8705 (hypergeometric test). Horizontal bars in violin plots indicate the median values.

### Modulation of titin mechanics by redox modifications

Our results show that evolutionary and structurally conserved cysteines of the I-band of titin are oxidized *in vivo*. We find reversible oxidations both in disulfide-competent cysteines and in unpaired cysteines that can establish S-thiolation adducts. These modifications have been proposed to induce opposite mechanical effects (*18, 21, 22*), an observation that may contribute to explain the different modulation of titin-based striated muscle stiffness under specific redox challenges (*22, 23, 29, 30*). To illustrate the range of regulation of titin mechanics by cysteine oxidation, we built on previous Monte Carlo simulations (*22*) to integrate all known mechanical effects of redox posttranslational modifications in titin Ig domains. These include reduction of contour length by disulfide bonds (*21*), higher unfolding and folding rates of disulfide-containing domains (*21, 41*), and higher unfolding rates and hampered folding of S-thiolated domains (*22*). In our simulations, we tuned unfolding and refolding rate constants to qualitatively reproduce the recently described unfolding/folding dynamics of native titin (*33*) (**Supplementary Note S1, Supplementary Figure S4A**). In the simulations, a virtual human I-band titin is subject to 1 Hz triangular force pulses between 0 and a predefined peak force, and the resulting length of titin is measured (**Figure 6A,B**). During the extension/relaxation cycles, titin domains unfold and refold stochastically according to their folding and unfolding rates, which are dependent on their redox state. At t = 0 s, all domains are folded and, as the simulations proceed, a fraction of domains transition to the unfolded state resulting in longer titin lengths at peak force (**Figure 6C,D**). The simulation time was long enough to reach steady-state lengths at all peak forces (**Figure 6C,D, Supplementary Figure S4B-E**).

**Figure 6.**
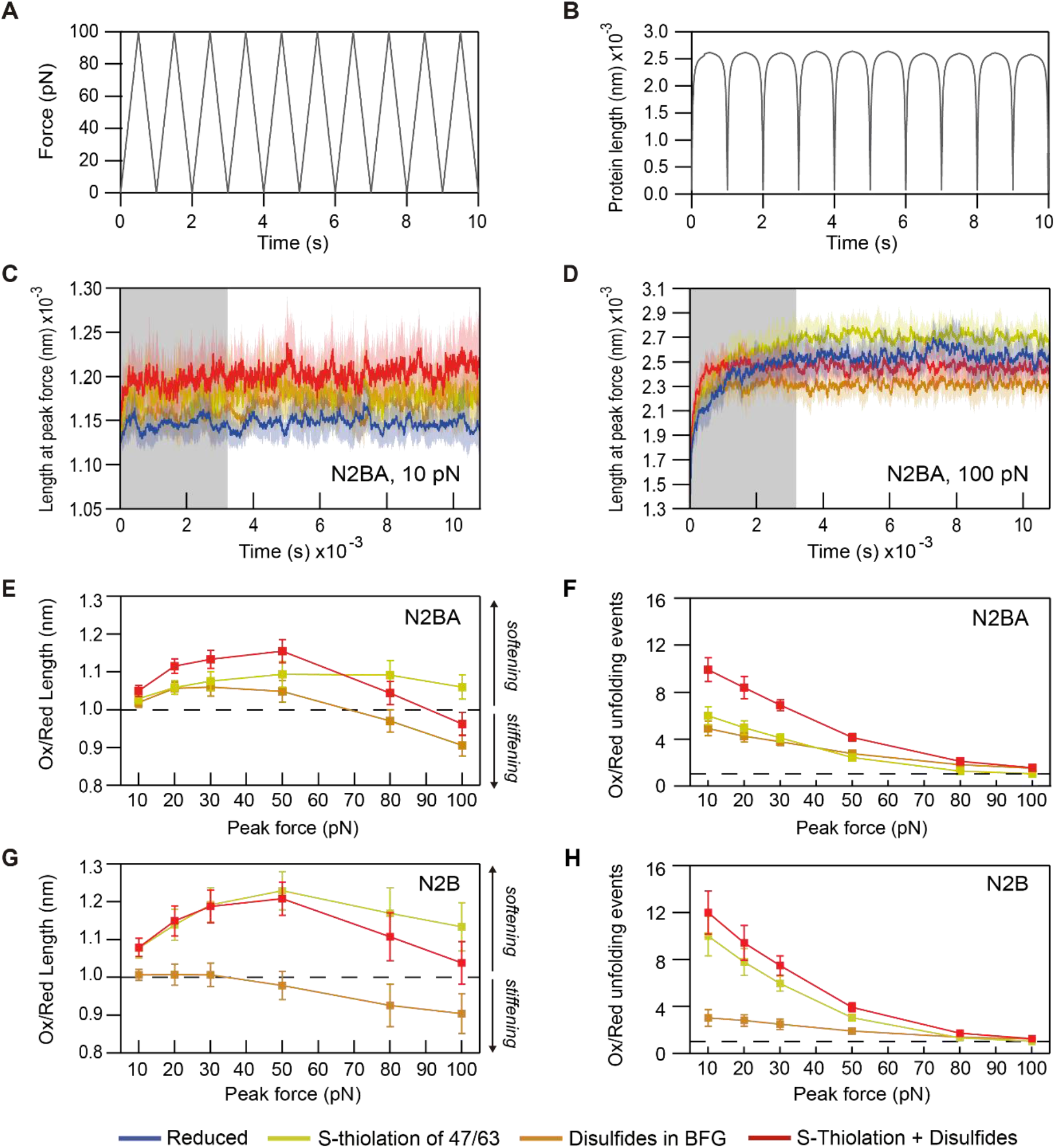
Effects of reversible cysteine oxidations on titin mechanics. **A**: Monte Carlo simulations subject virtual titins to a 1Hz oscillating triangular force pulse to a predefined peak force (100 pN in the example shown). **B**: Example of the length of reduced N2BA titin when pulled to a 100 pN peak force (10 seconds of simulation after reaching steady state are shown). **C,D**: Length of N2BA titin at peak force during simulations in which titin is reduced, or oxidized by disulfides, S-thiolation adducts or both (color code is indicated at the bottom; peak forces are indicated in the insets). Graphs show the average of 10 independent simulations and SD is indicated by shaded areas. For subsequent analyses, we considered times longer than 1 hour, in which the length of titin fluctuates around steady state values (region outside the grey shading). **E**: Ratio of oxidized *vs* reduced N2BA titin length at different peak forces. **F**: Ratio of oxidized *vs* reduced N2BA unfolding events in simulations at different peak forces. **G**: Ratio of oxidized *vs* reduced N2B titin length at different peak forces. **H**: Ratio of oxidized *vs* reduced N2B unfolding events in simulations at different peak forces. In panels E-H, n= 10 simulations; error bars: SD.

Simulations of the canonical N2BA titin at a low peak force of 10 pN show that disulfides and S-thiolations result in longer titin lengths (i.e. lower stiffness) (**Figure 6C**), while at a peak force of 100 pN, the effect of disulfides reverses leading to overall titin stiffening (**Figure 6D**). At this high peak force, S-thiolation maintains its softening effect. Additive mechanical modulation occurs if both oxidative modifications are present in titin simultaneously (**Figure 6C,D)**. To have a broader view of the extent of titin softening/stiffening induced by redox modifications, we did simulations at a range of peak forces and calculated the ratio of titin steady-state peak lengths between the oxidized and the reduced conditions. Results show that the softening effect of S-thiolation remains fairly constant, whereas at 50-80 pN peak force, the contribution of disulfides transitions from softening to stiffening (**Figure 6E, Supplementary Figure S4F**). This dual behavior stems from the fact that disulfides favor mechanical unfolding of Ig domains (softening effect), while also reducing the contour length of unfolded domains and increasing folding rates (stiffening effects). At low peak forces in which Ig domain unfolding rates are low, the softening effect is more prominent; while at high peak forces in which Ig unfolding is more frequent, the stiffening effects prevail. Beyond modulation of steady-state titin stiffness, our simulations also illustrate that both disulfides and S-thiolations induce a more dynamic state of titin by favoring Ig domain unfolding reactions, particularly at low forces (**Figure 6F, Supplementary Figure S4G**).

In the human heart, the short N2B isoform is expressed to similar levels as the longer and softer N2BA (*42, 43*). Since alternative splicing occurs at the region of titin with the highest density of cysteines (**Figure 1B**), we also ran Monte Carlo simulations for the N2B isoform of titin. In contrast to the results obtained with N2BA, we find that disulfides do not induce softening of N2B titin at any peak force (**Figure 6G, Supplementary Figure S4H-N**). Interestingly, S-thiolation softens N2B titin to a greater extent (20% *vs*. 10% for N2BA titin at 50 pN peak force, **Figure 6E,G**), reflecting the higher density of S-thiolation-competent Ig domains in N2B (**Supplementary Table S1**). For the same reason, the extent of modulation of titin dynamics by redox modifications is also different in N2B and N2BA titins (**Figure 6H, Supplementary Figure S4O**), although in both isoforms oxidations increase protein dynamics by favoring more Ig domain unfolding.

Our MS results were scarce with regards to the oxidation state of the cysteines in the N2Bus region if titin. Monte Carlo simulations show that potential disulfides in the N2Bus (*23*) would boost the overall stiffening effect of this redox modification, especially in the short N2B isoform (**Supplementary Figure S4P,Q**). Indeed, under conditions in which N2Bus cysteines form disulfides, disulfides always stiffen titin (**Supplementary Figure S4P,Q**). Taking together all the results from the Monte Carlo simulations, we conclude that reversible redox modifications have profound effects in the mechanics and dynamics of titin, in a manner that is highly dependent on the specific modification, the isoform of titin and the applied force protocol.

### Missense variants that affect conserved Cys3575 cause DCM

Having shown the functional relevance of titin conserved cysteines, we screened patients followed at the Heart Failure and Inherited Cardiac diseases Unit of Hospital Universitario Puerta de Hierro Majadahonda (Madrid, Spain) for variants targeting these residues. We identified a clear match in a patient who carried a Chr2(GRCh38):g.178741559A>T; p.Cys3575Ser variant (**Figure 7A**) (protein numbering according to Uniprot Q8WZ42). Cys3575, which is 100% conserved in 27 species ranging from zebrafish to human (**Supplementary Figure S5A)**, is the conserved cysteine F of the cardiac specific I21 domain (**Figure 7B**) and shows OS =1 both in human (**Figure 5C**) and mice (the equivalent position in mouse titin is Cys3581).

**Figure 7.**
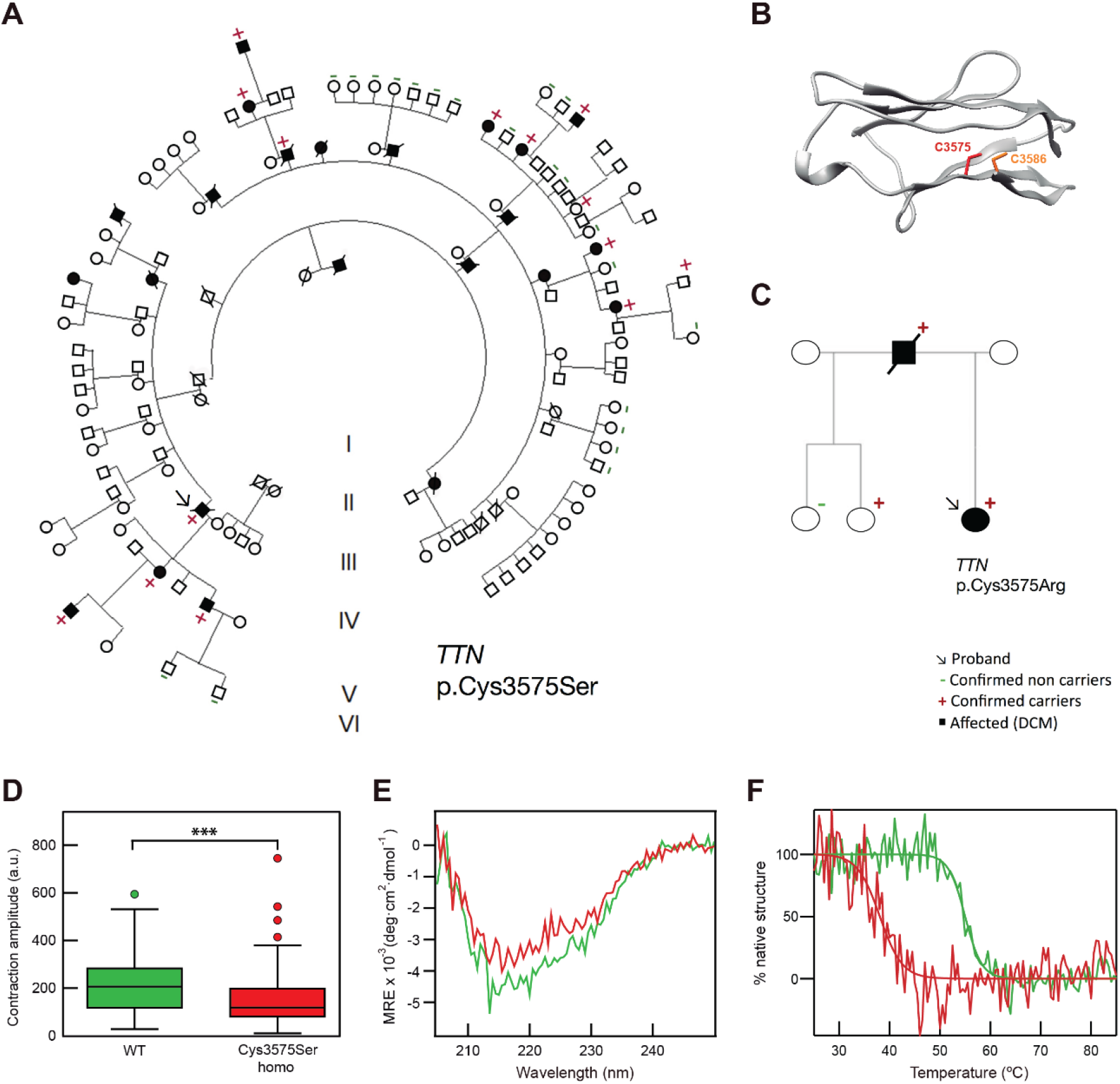
Mutations in conserved Cys3575 cause dilated cardiomyopathy. **A**: Pedigree of a large Spanish family with 14 TTN p.Cys3575Ser carriers (12 with DCM, penetrance 86%) and 22 non-carriers. **B**: Homology model of the I21 domain of titin, highlighting its two cysteine residues. **C**: Pedigree of a Danish family with the TTN p.Cys3575Arg variant, with 3 carriers (2 with DCM, penetrance 67%) and 1 non-carrier. **D**: Box-and-whiskers plots showing amplitude of contraction of WT (green, n = 87) and homozygous Cys3575Ser (red, n = 119) hiPSC-induced cardiomyocytes (p=0.0001, Mann-Whitney). Box encompasses data between quartiles 1 and 3 and horizontal lines represent the median of the distributions. Whiskers and outliers were calculated using Tukey’s method as implemented in Graph Pad. **E**: Far-UV circular dichroism spectra at 25°C of WT (green) and Cys3575Ser (red) recombinant I21 protein domains. **F**: Thermal unfolding curves of recombinant I21 WT (green) and Cys3575Ser (red) domains. Sigmoidal fits to the data are shown.

The proband was a male who underwent cardiac transplantation at 57 years of age and who carried no other pathogenic or likely pathogenic genetic variants in DCM-causing genes. 12/14 (86%) of the p.Cys3575Ser carriers in the family had DCM phenotype with a median age at DCM diagnosis of 33 years (interquartile range, IQR: 18-45) (**Figure 7A**). Mean left ventricular ejection fraction (LVEF) was 43 ± 6 % and mean left ventricular end-diastolic diameter (LVEDD) was 57 ± 4 mm, as determined by echocardiography (errors are SD). The two individuals who had the missense variant and did not show DCM phenotype were a 38 year-old man and a 64 year-old woman, although LVEF was in the lower limit of normal range in both cases (50-55%). A total of 22 additional relatives were non-carriers of the variant and all showed a normal phenotype (median age 51 years, IQR: 24-60). Family specific two-point logarithm of the odds (LOD) score was 3.96 using a dominant model and 80% penetrance, which strongly supports the linkage between phenotype and variant (*44*). Following this finding, we approached other European inherited cardiac disease units for genetically sequenced DCM patients with missense variants in Cys3575. An additional Danish DCM family captured our attention. In this family the same Cys3575 is mutated but in this case Cys is replaced by Arg (g.178741559A>G). Cosegregation analysis in this smaller family was limited as only two individuals showed DCM (**Figure 7C**). The proband was a young female with a very low LVEF (20%) and a severely dilated left ventricle (LVEDD = 75 mm) who underwent heart transplantation at 17 years of age. Her father was diagnosed with DCM when he was 54 years old and died at the age of 73 years. Genetic analyses of his kindred showed that the father was an obligate carrier of the p.Cys3575Arg variant supporting the pathogenic nature of the variant. The 52-year-old proband’s half-sister who also carries the variant had normal cardiac phenotype but a borderline depressed global longitudinal strain of −17%.

To provide functional evidence that the variants are pathogenic, we undertook cellular and molecular phenotyping studies. Human-induced-pluripotent-stem-cell (hiPSC)-derived cardiomyocytes carrying the p.Cys3575Ser variant in homozygosity show deficient contractility (**Figure 7D**), further supporting pathogenicity (*45*). Similarly to previous results (*45*), we did not detect any functional deficit in heterozygous p.Cys3575Ser cells (**Supplementary Figure S5B**). At the DNA level, the mutation lies in exon 49 of titin, more than 100 bp away from exon-exon junctions. Therefore, it is highly unlikely that the mutations interfere with native mRNA splicing sites, and the probability of inducing new splicing sites is also very low according to bioinformatics predictions (**Supplementary Figure S5C,D**). Using far-UV circular dichroism, we observed that a recombinant version of I21 Cys3575Ser domain (**Supplementary Note S2, Supplementary Figure S5E**) preserves the overall wild-type fold (**Figure 7E**) but is barely stable at physiological temperatures (melting temperature, T_m_ = 38 ± 1 °C *vs* 55 ± 1°C for wild-type domain, errors are SD of the sigmoidal fit, **Figure 7F, Supplementary Figure S5F**). Given the highly dissimilar physicochemical properties of Arg and Cys, we expect the Cys3575Arg mutation to lead also to strong domain destabilization.

Our MS results show that Cys3575 is oxidized in cardiac titin. The parent I21 domain contains a second highly conserved cysteine (residue 3586 in human titin), which is located in strand G at a distance that is compatible with disulfide bond formation with Cys3575 (*21*) (**Figure 7B**). The MS data were inconclusive with regards to the oxidation state of Cys3586 since mBBr-derivatized peptides were found for this position also in –DTT control samples. No mBBr-modified peptides for the equivalent Cys3592 in P0 mouse titin were detected. In both DCM variants targeting Cys3575, the disulfide between positions 3575 and 3586 cannot be established, which would leave the mutant domains in a highly destabilized state according to our circular dichroism data (**Figure 7F**).

In summary, our results show that Cys3575, which we demonstrate to be oxidized in human myocardium, is a target of DCM mutations that destabilize the parent domain and alter the mechanical function of hiPSC-derived cardiomyocytes.

## DISCUSSION

The high nucleophilicity of the thiol side chain makes cysteine the most reactive protein amino acid. In addition, cysteine residues are sensitive to irreversible oxidation, which can result in protein aggregation and degradation (*46*). As a consequence, proteins tend to include cysteine only if the associated functional benefits compensate the risks stemming from its peculiar physicochemical properties. Indeed, cysteine is among the least frequent but most conserved residues in proteins (*47*), reflecting key functional roles including configuration of enzyme active sites or metal chelation sites (*48, 49*), establishment of disulfide bonds (*41*), deployment of electron transport systems (*50*), or as redox sensors (*51*). Although several *in vitro* experiments over the last years have shown that cysteine oxidation is a strong modulator of titin mechanics, evidence that such modulation is functionally relevant *in vivo* has been lacking (*18*).

*In vitro*, conserved cysteines B, F and G can be induced to form disulfide bonds that alter the nanomechanics of the parent Ig domains (*21, 24*). Hence, a plausible scenario is that equivalent disulfides are present naturally in native titin as mechanical rheostats. This view has been traditionally considered unlikely taking into consideration the reducing environment of the cytosol in which sarcomeres are located (*23, 31*). However, our biochemical and MS results strongly support the existence of disulfides in titin in basal conditions in both human and mice (**Figure 2–5**), which adds to the increasing pool of evidence that redox compartmentalization of cardiomyocytes is complex (*52*). In this regard, future research will aim at identifying the biochemical systems responsible for disulfide formation in titin (*53–55*), and whether equivalent constitutive oxidations target other proteins in the sarcomere (*21, 56-58*).

Our results indicate that titin I-band cysteines that cannot establish disulfide bonds, such as conserved Cys47, can also be oxidized in basal conditions. By integrating current knowledge on the mechanical effects of cysteine oxidations obtained at the single-molecule level, our Monte Carlo simulations have allowed us to explore the range of oxidative mechanical modulation of titin (**Figure 6**). The simulations suggest that the extent and direction of mechanical modulation depends on the specific titin isoform, the biochemical nature of the oxidative modification (disulfide *vs* S-thiolation) and the range of forces experienced by titin. Two observations stemming from the Monte Carlo simulations are relevant for the redox modulation of titin mechanics at physiologically relevant forces, which are generally assumed to be < 10 pN/titin molecule (*19, 59–61*). First, due mainly to increased Ig unfolding rates, oxidations render the titin filament much more dynamic, up to one order of magnitude at low forces (**Figure 6F,H**). More frequent Ig unfolding can change the landscape of interactors, modulating titin-based mechanosignaling (*42*), whereas high folding rates enabled by disulfide bonds can sustain titin’s contribution to active muscle contraction (*33, 59, 62*). Monte Carlo simulations also illustrate that disulfides can result both in stiffening or softening of titin (**Figure 6E,G**), a consequence of the opposing effects of disulfides on contour length and the (un)folding rates of Ig domains. This observation, together with the softening effects of S-thiolation, can reconcile seemingly contradictory results of experiments using muscle preparations. For instance, treatment of human cardiomyocytes with the reducing enzyme thioredoxin results in drops in passive tension during oscillatory changes in length (*23*) while incubation with DTT results in increased passive tension during stepwise length increases (*22*). Similar DTT incubations also increase passive tension of rat and human skeletal fibers (*30*). Regarding oxidative modifications, treatment of mouse cardiac muscle with H2O2 induces increased passive tension in oscillatory protocols (*29*), whereas specific S-thiolation reactions lead to softening during stepwise extension of human cardiomyocytes (*22*), an effect also observed in human and rat skeletal muscle (*30*).

Overall, the Monte Carlo simulations show that the range of mechanical modulation of titin achieved by redox posttranslational modifications is ample. We have shown that right after birth, cardiac titin becomes more oxidized (**Figure 3**). We speculate that oxidation can allow titin to rapidly fulfil more demanding *ex utero* mechanical functions (*63*), even before the isoform content completes transition to an adult profile over a longer timescale of days (*4, 38*) (**Supplementary Figure S3B**). While under these physiological conditions the boost of titin oxidation is beneficial, we speculate that altered redox signaling may cause titin dysfunction during disease (*57*), for instance contributing to myocardial stiffening following myocardial infarction (*64*).

Our functional and conservation analyses show that cysteines are important for the modulation of titin mechanics, narrowing screening efforts to identify missense mutations that cause DCM. As a consequence, we were able to demonstrate that alteration of the conserved Cys3575, a residue which we detect to be oxidized in cardiac titin, is responsible for DCM in humans (**Figure 7**). Moreover, from a mechanistic point of view, we have detected extensive domain destabilization of the Cys3575Ser mutant, which may trigger excessive titin degradation and similar downstream effects as in the case of titin truncating variants. We have recently demonstrated a similar scenario in missense mutations of cardiac myosin-binding protein C that cause hypertrophic cardiomyopathy (*65*). We speculate that given the important role of cysteines in modulating titin mechanics, the fold of titin Ig domains may have evolved to accommodate these residues, and that missense mutations targeting conserved cysteines may result in strong destabilization and lead to disease.

The notion that missense mutations in titin could cause DCM is not new; however, our study is the first to unequivocally confirm that *TTN* missense variants can cause DCM in humans. In 2002, Gerull *et al*. reported that the missense mutation Trp976Arg in Ig domain 3 of titin (dbSNP: rs267607155), which was found in a moderately big DCM family, could cause the disease (*16*). The calculated LOD score for this family is 2.73 assuming an 80% penetrance of the disease, which is below the value of 3 usually considered to be associated with causal mutations (*44*). Interestingly, in our alignment, Trp976 is 100% evolutionary conserved along 35 species, and it is also 99% structurally conserved in human and mouse Ig domains highlighting its potential role in Ig domain stabilization (*16*). Since the identification of Trp976Arg, there have been anecdotal reports suggesting that *TTN* missense variants can be indeed pathogenic (*10, 66*). A spontaneous canine DCM model caused by a missense variant has been reported. This model exhibited a *TTN* missense variant in the Ig-like domain I71 in the I-band of the cardiac N2BA isoform that was significantly associated with DCM (p < 0.0001) in a family of Doberman pinschers. This variant corresponds to p.Gly8898Arg (ENST00000589042.5) in humans and it affects a 100% conserved glycine in our alignment (*67*). In humans, a 2015 paper reported four small DCM kindreds with 5 missense rare variants in *TTN* considered “severe” according to bioinformatic tools (*68*). As there were only 2 DCM individuals genotyped per family in the study, the evidence of pathogenicity was weak due to limited cosegregation and the case for *TTN* missense variants as definite cause of DCM remained open. Rare missense variants in *TTN* are extremely frequent hampering functional classification when interpreting NGS studies. This is exemplified by a recent study showing that in 530 DCM patients, almost 7% of them had rare *TTN* missense variants predicted to be deleterious by bioinformatics filtering. When compared to a large reference population database (ExAC) there was not a significant enrichment of the rare missense variants in the DCM population compared with ExAC and authors concluded that *TTN* missense variants should be classified as likely benign in the clinical diagnostic workflow (*17*). Our results confront with this conclusion and have direct application into clinical practice. Currently, rare *TTN* missense variants found in DCM genetic studies are considered unknown significance or likely benign and no further action is pursued. Based on our demonstration of the important biological role of titin cysteines and that missense mutations can cause DCM, now if a rare missense variant affecting a conserved cysteine in *TTN* is identified, predictive genetic testing might be performed. In these cases, functional assays and evaluation of the variant in relatives to confirm cosegregation should be undertaken. Similar strategies could be put in place in the future for other evolutionary and structurally conserved residues in titin.

## METHODS

### Human subject research

Human subject research was carried out in accordance with principles outlined in the Declaration of Helsinki. Probands with DCM were systematically studied by NGS with a panel of 121 genes associated or possibly associated with DCM at a certified clinical laboratory. These analyses did not show any pathogenic/likely pathogenic rare variants. Relatives were genetically studied for *TTN* missense variants found in probands by Sanger sequencing. Additionally, three phenotype-positive distant relatives of the family with the p.Cys3575Ser variant underwent exome sequencing that did not reveal commonly shared additional rare variants in cardiomyopathy-associated genes. Cardiac evaluation of all subjects included ECG and echocardiography. Selected individuals underwent cardiac magnetic resonance and additional cardiac tests according to clinical practice. DCM was defined as a left ventricular ejection fraction (LVEF) <50% (*69*). All individuals provided written consent. Study was approved by Hospital Universitario Puerta de Hierro ethics committee. Procedures for procurement of human left ventricular tissue from non-failing donor hearts that were not accepted for transplantation were approved by the Ethical Committee of the Medical University of Graz (28-508 ex 15/16) and the *Instituto de Salud Carlos III* (CEI PI 65_2017-v1). Upon ice-cold cardioplegia, cardiac biopsies were harvested from the left ventricular free wall, quickly frozen in liquid nitrogen and stored at −80°C. Non-failing hearts had echocardiographic evidence of preserved ejection fraction (>50%) and a clinical history that was free of cardiac abnormalities.

### Animal research

All animals used in this work were CD1 mice housed and maintained in the animal facility at the CNIC (Madrid, Spain) in accordance with national and European Legislation. Procedures were approved by the CNIC Animal Welfare Ethics Committee and by the Area of Animal Protection of the Regional Government of Madrid (PROEX 042/18).

### Cysteine conservation analysis

The longest currently available titin sequences for the 37 species reported in (*24*) were aligned using Clustal Omega, with the only exception of chicken, for which no titin sequence with enough coverage was found in Uniprot, GenomeNet or NCBI databases. Sequence identification codes are listed in **Supplementary Note S3**. Conservation values for all cysteine positions in the human titin sequence (Uniprot Q8WZ42) were obtained as percentages of conservation not considering gaps in the alignment. The vast majority of cysteine positions show very high occupancy (98% average occupancy, **Supplementary Figure S1**), which supports robustness of the conservation analyses. The alignment and homology models of titin Ig domains of human titin have been published before (*21*); alignment of mouse titin Ig domains appearing in Uniprot was obtained similarly using Clustal Omega (**Supplementary Note S4**). These Ig domain alignments were used to classify titin cysteines for structural conservation and mass spectrometry analyses.

### In-gel determination of reversibly oxidized thiols

Our protocol for in-gel determination of reversibly oxidized thiols was adapted and optimized from previous reports (*24, 32*). Protein extracts were obtained by cryopulverization of myocardial tissue followed by homogenization in sample buffer (50 mM Tris-HCl, 10 mM EDTA, 3% SDS, pH 6.8; 40 μl per mg of tissue) containing 50 mM NEM, unless indicated otherwise. Samples were run on SDS-PAGE gels (3.5% for analysis of titin, 12% for other proteins), and oxidized thiols were then reduced by incubation of the gel with 10 mM DL-dithiothreitol (DTT) (Sigma-Aldrich) in 50 mM ammonium bicarbonate, pH 8.8 at 60°C. After 3 washes of 20 min with sample buffer, the newly reduced thiols were labeled by incubation with 5 mM mBBr (Merk Millipore) in sample buffer during 2 h at room temperature in the dark. The excess of mBBr was removed by three washes with destaining solution (40% ethanol, 10% acetic acid) lasting 1 h, overnight and again 1 h. Fluorescent bands were visualized using a Gel-Doc (BioRad) with UV excitation (standard filters for ethidium bromide). Coomassie staining of the same gel was used to normalize fluorescence signals. Quantification of the bands was done by densitometry using Quantity One. Fluorescence signals coming from a replicate gel not treated with DTT were subtracted. Building on a previous report (*24*), control samples prepared by mixing different proportions of oxidized (I91–32/75)_8_ polyprotein and its Cys-free version were also included in the experiment in order to build calibration curves (**Supplementary Figure S3**). Two terminal cysteines in the polyproteins were not considered in these calculations since their oxidation status could not be ascertained. When comparing oxidation of different proteins, we normalized the oxidation signals by their density of cysteines (Uniprot entries Q8WZ42-1 and -3 for human N2BA and N2B respectively, A2ASS6-1 and -2 for mouse N2BA and N2B respectively, and Q02566 for mouse cardiac myosin).

### Mass spectrometry

Titin bands were sliced from SDS-PAGE gels used for fluorescence quantifications, diced and washed by incubation with 200 μl high purity water (Fluka CHROMASOLV™ LC-MS) for 10 min at 1200 rpm (4 times). The resulting gel pieces were dehydrated by 2 incubations with 100% acetonitrile and 1 incubation with 50 mM ammonium bicarbonate (pH 8.8) in acetonitrile (both incubations, 15 min shaking at 1200 rpm). After drying using a SpeedVac, the gel pieces were incubated with modified trypsin from porcine pancreas (Sigma Aldrich) in digestion buffer (10% acetonitrile in 50 mM ammonium bicarbonate, pH 8.8) for 2 h in ice to allow the diffusion of the inactive enzyme, and then at 37°C overnight. Digested tryptic peptides were extracted with 1% trifluoroacetic acid in acetonitrile (15 min incubation, 1200 rpm shaking) and dried. Finally, they were resuspended in 1% trifluoroacetic acid by pulse-vortexing and sonication, and desalted using OMIX commercial columns (Biomaster group). Peptides were then injected into a reversed phase C-18 nano-column (Acclaim PepMap RSLC, 75 μm internal diameter and 50 cm length), and eluted to be analysed in a hybrid quadrupole-Orbitrap Q Exactive mass spectrometer (Thermo Scientific) for protein identification. A continuous acetonitrile gradient consisting of 0-30% A for 120 min, 50-90% B for 3 min (A= 0.1% formic acid; B= 98% acetonitrile, 0.1% formic acid) at a flow rate of 200 nL/min was used to elute tryptic peptides from the nano-column to a nanospray emitter for real time ionization and induced fragmentation. High resolution mass spectra were acquired in a data-dependent manner with dynamic exclusion by combining a MS spectrum (from 400-1500 m/z, 120,000 resolution) followed by the MS/MS spectra (60,000 resolution) from the 15 most intense species. For protein identification, tandem mass spectra were extracted and charge state was deconvoluted by Proteome Discoverer 1.4.0.288 (Thermo Fisher Scientific). All MS/MS spectra were analyzed using SEQUEST (Thermo Fisher Scientific). Full proteome search databases (mouse: UniProtKB/Swiss-Prot April27_2016, 48736 sequences; human: UniProtKB/Swiss-Prot, November 2019, 74333 sequences) were supplemented with 116 cRAP proteins (common Repository of Adventitious Proteins, Global Proteome Machine). SEQUEST searches allowed two missed cleavages and used 20 ppm and 20 mDa precursor and fragment mass tolerances, respectively. NEM- or mBBr-modified cysteines, and oxidation of methionine were specified as variable modifications. In these initial searches, a target-decoy fixed value PSM validation strategy was used to filter peptides according to XCorr values (more than 1.2 in doubly-charged peptides, or more than 1.4 in triply-charged peptides for <5% FDR, and more than 1.42 in doubly-charged peptides, or more than 1.79 for triply-charged peptides for <1% FDR). Since the vast majority of the identified peptides belonged to titin in searches against full proteome databases, to optimize coverage of cysteine peptides we repeated the search using titin-only reduced search databases (*39*). Peptide identifications were validated using Vseq, an in-house developed tool that evaluates the mass tolerance, intensity, fragmentation goodness and quantitative value of MS/MS scans (*40*). Maximum matched ions value for a sequence was calculated considering the number of residues multiplied by 2, to cover the complete theoretical length of the main fragmentation series B and Y. Since experiments were acquired using a label-free strategy and using HCD for ion-induced dissociation, we set a threshold of 25% minimum matched ions, which corresponds to at least 50% of the Y-HCD-enhanced fragmentation series. We also filtered peptide identifications according to their E-scores, which is defined as the dot product of the intensities for all matched ions regardless of their charge state. We applied an E-score threshold of > 0.01. We verified that none of the unidentified scans showing a very low XCorr and matched ions value was above this limit. To estimate final FDR values, we repeated the searches considering the same parameters against the corresponding decoy databases. FDR was below 2% in all cases. To minimize false positive detection of oxidized cysteines, we analyzed in parallel the same samples without including DTT in the derivatization reactions. For each sample, cysteine positions belonging to mBBr-peptides identified both in the −DTT and +DTT treated specimens were excluded from the analysis (27 ± 5% and 30 ± 4% of the oxidized cysteine-containing peptides identified in mouse and in human). OS were calculated for each cysteine following

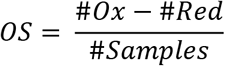

where #*Ox* and #*Red* are the number of samples in which the cysteine is detected as always oxidized or reduced, respectively, and #Samples is the total number of samples in which the cysteine is detected, including those instances in which both the oxidized and reduced forms are detected. OS ranges from −1 to 1; these extreme values are given to cysteines that are always detected as reduced or oxidized, respectively.

### Monte Carlo simulations

We constructed a simplified model of titin’s I band to predict the length of the molecule under force upon redox modifications using Monte Carlo simulations (*21, 22*). Simulations calculate the length of the two entropic regions of the protein (PEVK and N2Bus) and the Ig domains, according to the Freely Jointed Chain model of polymer elasticity and the Bell’s model of force-dependent reactions (**Supplementary Note S1**). The length of the entropic regions and the number of Ig domains in each isoform were determined according to human titin sequence in Uniprot, further curated in (*21*). Titin domains were classified according to their cysteine content, which was used to determine the modifications they could be target of (**Supplementary Table S1**) and their mechanical and kinetic parameters (**Supplementary tables S2-S5**). The constructed model was subject to an oscillating force protocol (triangular waves, 1 Hz, 10 ms simulation step) for 3 hours. Some simulations were also conducted using 1 ms simulation steps, leading to equivalent results. In the initial state all domains are folded. If oxidative modifications are present, they occur at all potential target sites. In conditions involving disulfide bonds, all domains capable of disulfide isomerization contain the BG disulfide in the initial state (*21*). We averaged 10 independent simulations for each condition. The code for the simulations and the subsequent analysis of the data was conducted in Igor Pro.

### hiPSC-derived cardiomyocytes characterization

*TTN* ^*Cys3575Ser*^-hiPSC lines were generated by using the CRISPR/Cas9 technology to introduce the corresponding point mutation in the TTN allele of a wild-type (WT) hiPSC line (HDF-iPS-SV10 - Spanish National Stem Cell Bank, ISCIII). The crRNA 5’-TGTATGGCCAGTAATGACTA-3’, was designed by the online CRISPOR-TEFOR software (http://crispor.tefor.net/crispor.py), matching the nucleotide involved in the mutation as the first base of sequence. The ssODN sequence lies in the No-PAM strand CTGTTTCAGTGTCTTTGTGACCCTCTCCTTTGGAATTAATTTTTAGATAGGCACTACATATTGTCTTTCC*G*TA*A*TC*G*TT<u>*G*CT*A*GC</u>CATAC**T***G*GT*G*TACTCTCCCTCATCCTCCAATTTGGTGAACAGAATGATCAGGCTATG, and was designed by inserting the intended mutation (**bold**), as well as silent variants (*italic)* to avoid unwanted cutting of the ribonucleoprotein complex (RPN) and to allow genotyping by restriction enzyme digestion (*Nhe*I-underlined). The primers used to genotype selected clones were hTTN-Fwd: 5’-CAGGTGGACTCACCCAACAT-3’ and hTTN-Rev 5’-CCAGGAATAAAGCAGAAGAAGGC-3’. Synthetic crRNA and tracrRNA (AltR CRISPRCas9 system, IDT) were mixed in equimolar concentrations (22 pmol/each), combined with 25 μM Cas9 Nuclease 3NLS to form the ribonucleoprotein complex. The RNP complex plus ssODN and AltR Cas9 Electroporator Enhancer (22 pmol/each) were electroporated, and cells were recovered and split into p100 dishes as described (*70*). After 7-10 days, colonies were picked up and split into p96 and p24 well plates for screening and maintenance respectively. DNA from 200 clones was isolated, PCR amplified and *Nhe*I digested. Several clones were identified as positive and Sanger sequenced. One proved to be homozygous and another one heterozygous for the desired mutation. For cardiac differentiation, WT and *TTN* ^*Cys3575Ser*^-hiPSC lines were expanded and dissociated into single cells as described (*70*). Briefly cells were seeded at a density of 1-1.2×10^6^ cells per well in Essential E8 medium supplemented with 10 μM ROCK inhibitor. Upon achieving confluence, they were treated with 8 μM GSK3 inhibitor (CHIR99021, Stemgent) in RPMI supplemented with B27 lacking insulin (Thermo Fisher). 24 hrs later, the medium was changed to RPMI/B27-insulin. On day 3, cells were treated with 5μM Wnt inhibitor IWP4 (Stemgent) in RPMI/B27 minus insulin medium, which was replaced two days later. From day 7 onwards, differentiated cells were maintained in RPMI/B27 medium supplemented with insulin. Beating areas were visible between days 8-10 from the onset of the differentiation. For iPSC-derived cardiac monolayer maturation, cells were disaggregated at day 32 by incubation with 0.25% trypsin-EDTA for 10 min at 37 °C and seeded onto matrigel-coated PDMS membranes (*71*) at a density of 1.25×10^5^ cells/well in 12-well plates. WT and *TTN* ^*Cys3575Ser*^-hiPSC derived cardiomyocytes (hiPSC-CMs) were maintained in RPMI/B27 medium supplemented with insulin for 1 week for further maturation. Video recording was taken and analyzed for single cell contraction amplitude using the Musclemotion algorithm (*72*).

### Protein purification

The *E. coli*-codon-optimized-cDNA coding for I21 WT was ordered to GeneArt and the Cys3575Ser mutant was produced by PCR-mutagenesis (sequences available in **Supplementary Note S2**). cDNAs were cloned in a custom-modified pQE80 expression plasmid (Qiagen) using BamHI and BglII restriction enzymes. Final expression plasmids were verified by Sanger sequencing. Domains were expressed in *E. coli* BLR(DE3). For the I21 WT domain cultures at OD_600_ = 0.6-1 were induced with 1 mM isopropyl β-D-1-thiogalactopyranoside (IPTG) and incubated for 3 h at 37°C. For the Cys3575Ser mutant, the culture was induced with 0.4 mM IPTG and incubated at 16°C overnight. Purification of His-tagged domains was achieved by metal affinity (using 1 mM DTT containing buffers) and gel filtration chromatographies following published protocols (*73*). Proteins were eluted from the final size-exclusion chromatography in 20 mM NaPi, pH 6.5, 63.6 mM NaCl and stored at 4°C. Purity of the preparations was evaluated by SDS-PAGE (**Supplementary Figure S5E**).

### Circular dichroism

CD spectra were collected using a Jasco J-810 spectropolarimeter. Purified proteins were tested in 20 mM NaPi, pH 6.5 and 63.6 mM NaCl at 0.3 mg/ml protein concentration in 0.1-cm-pathlength quartz cuvettes. Protein concentration was obtained from A_280_ values using theoretical extinction coefficients estimated using ProtParam (*74*) (E^0.1%^ = 0.925). Spectra were recorded at 50 nm/min scanning speed and a data pitch of 0.2 nm. Four scans were averaged to obtain the final spectra. The contribution of the buffer was subtracted and spectra were normalized by peptide bond concentration. To study thermal denaturation, CD signal at 215 nm was monitored as temperature increased from 25 to 85 °C at a rate of 30°C/h. Temperature control was achieved using a Peltier thermoelectric system. To estimate T_m_, changes in CD signal were fit to a sigmoidal function using IGOR Pro (Wavemetrics).

### Statistics

Statistical tests were run using Graph Pad Prism 8.3.1 for Windows. To evaluate the tendency of different categories of cysteines to be oxidized (OS>0), we estimated the corresponding p-values using the hypergeometric distribution function implemented in Excel (Microsoft). Errors are given by SEM, unless indicated otherwise.

## Acknowledgements

We thank CNIC’s Pluripotent Cell Technology Unit for excellent technical support.

## Funding

This work was supported by the *Ministerio de Ciencia e Innovación* grants BIO2014-54768-P, BIO2017-83640-P, and RYC-2014-16604 to JAC, and RTI2018-096961-B-I00 to ELP, the Regional Government of Madrid grants S2018/NMT-4443 and PEJ16/MED/TL-1593 to JAC, the *Instituto de Salud Carlos III* (ISCIII) grants PI17/01941 to PGP and CPII14/00027 to ELP, and by a Mutual Medical Research Award 2017 to FD. We acknowledge funding from the European Research Area Network on Cardiovascular Disease through grants Genprovic to PGP (ISCIII-AC16/0014) and MINOTAUR to SS (The Austrian Science Fund – FWF, I3301-B31) and JAC (ISCIII-AC16/00045). We acknowledge support from the RD012/0042/0066 ISCIII grant to PGP and ELP. The CNIC is supported by ISCIII, the *Ministerio de Ciencia e Innovación* and the Pro CNIC Foundation, and is a Severo Ochoa Center of Excellence (SEV-2015-0505). IMM was the recipient of a CNIC-ACCIONA Masters Fellowship. CSC is the recipient of an FPI-SO predoctoral fellowship BES-2016-076638.

## Author contributions

PGP and JAC conceived and led the project. EHG, IMM, CSC, MRP, DG and RPJ analyzed titin sequence. EHG, IMM, CSG, NV and CBC did in-gel determination of reversible cysteine oxidations. EHG, IMM, EBK, EC and JV did and analyzed mass spectrometry experiments. MA, SS, and PPR procured and characterized human heart samples. EHG, IMM, NV and DVC cloned, purified and analyzed recombinant proteins. EHG, AFT, JV and JAC did statistical analysis of experimental data. IMM did Monte Carlo simulations. FD, EGL, MCM, BB, AB, JPO, JMGA TBR, TMH, IVO, JPF and PGP obtained and interpreted clinical and genetics data. LL, BP, FGA, GG, JAB and ELP obtained hiPSC data. EHG, FD, IMM, PGP and JAC drafted the manuscript with input from all authors.

## Competing interests

The authors declare no competing interests.

## Data and material availability

Data and materials are available on reasonable request to the corresponding authors. Mass spectrometry raw data have been deposited in PeptideAtlas (Identifier: PASS01609).

## List of Supplementary Materials

Supplementary File S1,

Supplementary Figures S1-S5,

Supplementary Notes S1-S4,

Supplementary Tables S1-S5

References (75-77)

## Notes

### Competing Interest Statement

The authors have declared no competing interest.

## References and Notes

1. J. K. Freundt, W. A. Linke, Titin as a force-generating muscle protein under regulatory control. J Appl Physiol (1985) 126, 1474–1482 (2019).

2. B. R. Anderson, H. L. Granzier, Titin-based tension in the cardiac sarcomere: Molecular origin and physiological adaptations. Progress in Biophysics and Molecular Biology 110, 204–217 (2012).

3. E. C. Eckels, R. Tapia-Rojo, J. A. Rivas-Pardo, J. M. Fernandez, The Work of Titin Protein Folding as a Major Driver in Muscle Contraction. Annual review of physiology 80, 327–351 (2018).

4. A. Opitz Christiane, C. Leake Mark, I. Makarenko, V. Benes, A. Linke Wolfgang, Developmentally Regulated Switching of Titin Size Alters Myofibrillar Stiffness in the Perinatal Heart. Circulation Research 94, 967–975 (2004).

5. A. Freiburg, K. Trombitas, W. Hell, O. Cazorla, F. Fougerousse, T. Centner, B. Kolmerer, C. Witt, J. S. Beckmann, C. C. Gregorio, H. Granzier, S. Labeit, Series of exon-skipping events in the elastic spring region of titin as the structural basis for myofibrillar elastic diversity. Circulation research 86, 1114–1121 (2000).

6. M. Kruger, in Cardiac Cytoarchitecture, E. Ehler, Ed. (Springer International Publishing, 2015), pp. 109–124.

7. D. S. Herman, L. Lam, M. R. G. Taylor, L. B. Wang, P. Teekakirikul, D. Christodoulou, L. Conner, S. R. DePalma, B. McDonough, E. Sparks, D. L. Teodorescu, A. L. Cirino, N. R. Banner, D. J. Pennell, S. Graw, M. Merlo, A. Di Lenarda, G. Sinagra, J. M. Bos, M. J. Ackerman, R. N. Mitchell, C. E. Murry, N. K. Lakdawala, C. Y. Ho, P. J. R. Barton, S. A. Cook, L. Mestroni, J. G. Seidman, C. E. Seidman, Truncations of Titin Causing Dilated Cardiomyopathy. New Engl J Med 366, 619–628 (2012).

8. D. Kellermayer, J. E. Smith, H. Granzier, Titin mutations and muscle disease. Pflügers Archiv - European Journal of Physiology 471, 673–682 (2019).

9. D. Fatkin, I. G. Huttner, J. C. Kovacic, J. G. Seidman, C. E. Seidman, Precision Medicine in the Management of Dilated Cardiomyopathy. Journal of the American College of Cardiology 74, 2921 (2019).

10. J. S. Ware, S. A. Cook, Role of titin in cardiomyopathy: from DNA variants to patient stratification. Nature Reviews Cardiology 15, 241–252 (2018).

11. S. Cuenca, M. J. Ruiz-Cano, J. R. Gimeno-Blanes, A. Jurado, C. Salas, I. Gomez-Diaz, L. Padron-Barthe, J. J. Grillo, C. Vilches, J. Segovia, D. Pascual-Figal, E. Lara-Pezzi, L. Monserrat, L. Alonso-Pulpon, P. Garcia-Pavia, Genetic basis of familial dilated cardiomyopathy patients undergoing heart transplantation. The Journal of Heart and Lung Transplantation 35, 625–635 (2016).

12. S. van Heesch, F. Witte, V. Schneider-Lunitz, J. F. Schulz, E. Adami, A. B. Faber, M. Kirchner, H. Maatz, S. Blachut, C. L. Sandmann, M. Kanda, C. L. Worth, S. Schafer, L. Calviello, R. Merriott, G. Patone, O. Hummel, E. Wyler, B. Obermayer, M. B. Mucke, E. L. Lindberg, F. Trnka, S. Memczak, M. Schilling, L. E. Felkin, P. J. R. Barton, N. M. Quaife, K. Vanezis, S. Diecke, M. Mukai, N. Mah, S. J. Oh, A. Kurtz, C. Schramm, D. Schwinge, M. Sebode, M. Harakalova, F. W. Asselbergs, A. Vink, R. A. de Weger, S. Viswanathan, A. A. Widjaja, A. Gartner-Rommel, H. Milting, C. Dos Remedios, C. Knosalla, P. Mertins, M. Landthaler, M. Vingron, W. A. Linke, J. G. Seidman, C. E. Seidman, N. Rajewsky, U. Ohler, S. A. Cook, N. Hubner, The Translational Landscape of the Human Heart. Cell 178, 242–260 e229 (2019).

13. A. M. Roberts, J. S. Ware, D. S. Herman, S. Schafer, J. Baksi, A. G. Bick, R. J. Buchan, R. Walsh, S. John, S. Wilkinson, F. Mazzarotto, L. E. Felkin, S. Gong, J. A. MacArthur, F. Cunningham, J. Flannick, S. B. Gabriel, D. M. Altshuler, P. S. Macdonald, M. Heinig, A. M. Keogh, C. S. Hayward, N. R. Banner, D. J. Pennell, D. P. O’Regan, T. R. San, A. de Marvao, T. J. Dawes, A. Gulati, E. J. Birks, M. H. Yacoub, M. Radke, M. Gotthardt, J. G. Wilson, C. J. O’Donnell, S. K. Prasad, P. J. Barton, D. Fatkin, N. Hubner, J. G. Seidman, C. E. Seidman, S. A. Cook, Integrated allelic, transcriptional, and phenomic dissection of the cardiac effects of titin truncations in health and disease. Science translational medicine 7, 270ra276 (2015).

14. C. A. Tharp, M. E. Haywood, O. Sbaizero, M. R. G. Taylor, L. Mestroni, The Giant Protein Titin’s Role in Cardiomyopathy: Genetic, Transcriptional, and Post-translational Modifications of TTN and Their Contribution to Cardiac Disease. Frontiers in Physiology 10, (2019).

15. M. Gigli, R. L. Begay, G. Morea, S. L. Graw, G. Sinagra, M. R. G. Taylor, H. Granzier, L. Mestroni, A Review of the Giant Protein Titin in Clinical Molecular Diagnostics of Cardiomyopathies. Frontiers in Cardiovascular Medicine 3, (2016).

16. B. Gerull, M. Gramlich, J. Atherton, M. McNabb, K. Trombitas, S. Sasse-Klaassen, J. G. Seidman, C. Seidman, H. Granzier, S. Labeit, M. Frenneaux, L. Thierfelder, Mutations of TTN, encoding the giant muscle filament titin, cause familial dilated cardiomyopathy. Nat Genet 30, 201–204 (2002).

17. O. Akinrinade, T. Heliö, R. H. Lekanne Deprez, J. D. H. Jongbloed, L. G. Boven, M. P. van den Berg, Y. M. Pinto, T.-P. Alastalo, S. Myllykangas, K. v. Spaendonck-Zwarts, J. P. v. Tintelen, P. A. van der Zwaag, J. Koskenvuo, Relevance of Titin Missense and Non-Frameshifting Insertions/Deletions Variants in Dilated Cardiomyopathy. Scientific Reports 9, 4093 (2019).

18. E. Herrero-Galán, I. Martínez-Martín, J. Alegre-Cebollada, Redox regulation of protein nanomechanics in health and disease: Lessons from titin. Redox Biology 21, 101074 (2019).

19. H. Li, W. A. Linke, A. F. Oberhauser, M. Carrion-Vazquez, J. G. Kerkvliet, H. Lu, P. E. Marszalek, J. M. Fernandez, Reverse engineering of the giant muscle protein titin. Nature 418, 998–1002 (2002).

20. M. S. Kellermayer, L. Grama, Stretching and visualizing titin molecules: combining structure, dynamics and mechanics. J Muscle Res Cell Motil 23, 499–511 (2002).

21. D. Giganti, K. Yan, C. L. Badilla, J. M. Fernandez, J. Alegre-Cebollada, Disulfide isomerization reactions in titin immunoglobulin domains enable a mode of protein elasticity. Nature communications 9, 185 (2018).

22. J. Alegre-Cebollada, P. Kosuri, D. Giganti, E. Eckels, J. A. Rivas-Pardo, N. Hamdani, C. M. Warren, R. J. Solaro, W. A. Linke, J. M. Fernandez, S-glutathionylation of cryptic cysteines enhances titin elasticity by blocking protein folding. Cell 156, 1235–1246 (2014).

23. A. Grutzner, S. Garcia-Manyes, S. Kotter, C. L. Badilla, J. M. Fernandez, W. A. Linke, Modulation of titin-based stiffness by disulfide bonding in the cardiac titin N2-B unique sequence. Biophys J 97, 825–834 (2009).

24. A. Manteca, J. Schonfelder, A. Alonso-Caballero, M. J. Fertin, N. Barruetabena, B. F. Faria, E. Herrero-Galan, J. Alegre-Cebollada, D. De Sancho, R. Perez-Jimenez, Mechanochemical evolution of the giant muscle protein titin as inferred from resurrected proteins. Nature structural & molecular biology 24, 652–657 (2017).

25. B. S. Avner, K. M. Shioura, S. B. Scruggs, M. Grachoff, D. L. Geenen, D. L. Helseth, M. Farjah, P. H. Goldspink, R. J. Solaro, Myocardial infarction in mice alters sarcomeric function via post-translational protein modification. Mol Cell Biochem 363, 203–215 (2012).

26. B. Bodi, E. P. Toth, L. Nagy, A. Toth, L. Martha, A. Kovacs, G. Balla, T. Kovacs, Z. Papp, Titin isoforms are increasingly protected against oxidative modifications in developing rat cardiomyocytes. Free Radic Biol Med 113, 224–235 (2017).

27. J. Paulech, N. Solis, S. J. Cordwell, Characterization of reaction conditions providing rapid and specific cysteine alkylation for peptide-based mass spectrometry. Biochimica et Biophysica Acta (BBA) - Proteins and Proteomics 1834, 372–379 (2013).

28. H. Xiao, M. P. Jedrychowski, D. K. Schweppe, E. L. Huttlin, Q. Yu, D. E. Heppner, J. Li, J. Long, E. L. Mills, J. Szpyt, Z. He, G. Du, R. Garrity, A. Reddy, L. P. Vaites, J. A. Paulo, T. Zhang, N. S. Gray, S. P. Gygi, E. T. Chouchani, A Quantitative Tissue-Specific Landscape of Protein Redox Regulation during Aging. Cell 180, 968–983.e924 (2020).

29. J. Nedrud, S. Labeit, M. Gotthardt, H. Granzier, Mechanics on Myocardium Deficient in the N2B Region of Titin: The Cardiac-Unique Spring Element Improves Efficiency of the Cardiac Cycle. Biophysical Journal 101, 1385–1392 (2011).

30. D. Watanabe, C. R. Lamboley, G. D. Lamb, Effects of S-glutathionylation on the passive force–length relationship in skeletal muscle fibres of rats and humans. Journal of Muscle Research and Cell Motility, (2019).

31. O. Mayans, J. Wuerges, S. Canela, M. Gautel, M. Wilmanns, Structural evidence for a possible role of reversible disulphide bridge formation in the elasticity of the muscle protein titin. Structure 9, 331–340 (2001).

32. L. K. Rogers, B. L. Leinweber, C. V. Smith, Detection of reversible protein thiol modifications in tissues. Anal Biochem 358, 171–184 (2006).

33. J. A. Rivas-Pardo, Y. Li, Z. Mártonfalvi, R. Tapia-Rojo, A. Unger, Á. Fernández-Trasancos, E. Herrero-Galán, D. Velázquez-Carreras, J. M. Fernández, W. A. Linke, J. Alegre-Cebollada, A HaloTag-TEV genetic cassette for mechanical phenotyping of proteins from tissues. Nature communications 11, 2060 (2020).

34. K. Wang, J. McClure, A. Tu, Titin: major myofibrillar components of striated muscle. Proceedings of the National Academy of Sciences 76, 3698 (1979).

35. R. E. Hansen, J. R. Winther, An introduction to methods for analyzing thiols and disulfides: Reactions, reagents, and practical considerations. Analytical Biochemistry 394, 147–158 (2009).

36. J. Vonderen, J. J. van Vonderen, A. A. W. Roest, M. L. Siew, F. J. Walther, S. B. Hooper, A. B. te Pas, Measuring Physiological Changes during the Transition to Life after Birth. Neonatology 105, 230–242 (2014).

37. B. N. Puente, W. Kimura, S. A. Muralidhar, J. Moon, J. F. Amatruda, K. L. Phelps, D. Grinsfelder, B. A. Rothermel, R. Chen, J. A. Garcia, C. X. Santos, S. Thet, E. Mori, M. T. Kinter, P. M. Rindler, S. Zacchigna, S. Mukherjee, D. J. Chen, A. I. Mahmoud, M. Giacca, P. S. Rabinovitch, A. Aroumougame, A. M. Shah, L. I. Szweda, H. A. Sadek, The oxygen-rich postnatal environment induces cardiomyocyte cell-cycle arrest through DNA damage response. Cell 157, 565–579 (2014).

38. S. Lahmers, Y. Wu, D. R. Call, S. Labeit, H. Granzier, Developmental control of titin isoform expression and passive stiffness in fetal and neonatal myocardium. Circulation research 94, 505–513 (2004).

39. W. S. Noble, Mass spectrometrists should search only for peptides they care about. Nature Methods 12, 605–608 (2015).

40. S. Cogliati, E. Calvo, M. Loureiro, A. M. Guaras, R. Nieto-Arellano, C. Garcia-Poyatos, I. Ezkurdia, N. Mercader, J. Vázquez, J. A. Enriquez, Mechanism of super-assembly of respiratory complexes III and IV. Nature 539, 579–582 (2016).

41. P. Kosuri, J. Alegre-Cebollada, J. Feng, A. Kaplan, A. Ingles-Prieto, C. L. Badilla, B. R. Stockwell, J. M. Sanchez-Ruiz, A. Holmgren, J. M. Fernandez, Protein folding drives disulfide formation. Cell 151, 794–806 (2012).

42. W. A. Linke, N. Hamdani, Gigantic business: titin properties and function through thick and thin. Circulation research 114, 1052–1068 (2014).

43. O. Cazorla, A. Freiburg, M. Helmes, T. Centner, M. McNabb, Y. Wu, K. Trombitas, S. Labeit, H. Granzier, Differential expression of cardiac titin isoforms and modulation of cellular stiffness. Circ Res 86, 59–67 (2000).

44. D. R. Nyholt, All LODs Are Not Created Equal**A Microsoft Excel spreadsheet, for performing easy calculations of P values for the LOD scores described in this review, is available on request from the author. The American Journal of Human Genetics 67, 282–288 (2000).

45. J. T. Hinson, A. Chopra, N. Nafissi, W. J. Polacheck, C. C. Benson, S. Swist, J. Gorham, L. Yang, S. Schafer, C. C. Sheng, A. Haghighi, J. Homsy, N. Hubner, G. Church, S. A. Cook, W. A. Linke, S. Chen, J. G. Seidman, C. E. Seidman, HEART DISEASE. Titin mutations in iPS cells define sarcomere insufficiency as a cause of dilated cardiomyopathy. Science 349, 982–986 (2015).

46. M. Pajares, N. Jiménez-Moreno, I. H. K. Dias, B. Debelec, M. Vucetic, K. E. Fladmark, H. Basaga, S. Ribaric, I. Milisav, A. Cuadrado, Redox control of protein degradation. Redox Biology 6, 409–420 (2015).

47. S. M. Marino, V. N. Gladyshev, Cysteine Function Governs Its Conservation and Degeneration and Restricts Its Utilization on Protein Surfaces. Journal of Molecular Biology 404, 902–916 (2010).

48. S. Verma, R. Dixit, K. C. Pandey, Cysteine Proteases: Modes of Activation and Future Prospects as Pharmacological Targets. Frontiers in Pharmacology 7, (2016).

49. N. M. Giles, A. B. Watts, G. I. Giles, F. H. Fry, J. A. Littlechild, C. Jacob, Metal and Redox Modulation of Cysteine Protein Function. Chemistry & Biology 10, 677–693 (2003).

50. G. Bertoli, T. Simmen, T. Anelli, S. N. Molteni, R. Fesce, R. Sitia, Two Conserved Cysteine Triads in Human Ero1α Cooperate for Efficient Disulfide Bond Formation in the Endoplasmic Reticulum. Journal of Biological Chemistry 279, 30047–30052 (2004).

51. C. Klomsiri, P. A. Karplus, L. B. Poole, Cysteine-Based Redox Switches in Enzymes. Antioxidants & Redox Signaling 14, 1065–1077 (2010).

52. L. Swain, A. Kesemeyer, S. Meyer-Roxlau, C. Vettel, A. Zieseniss, A. Güntsch, A. Jatho, A. Becker, S. Nanadikar Maithily, B. Morgan, S. Dennerlein, M. Shah Ajay, A. El-Armouche, O. Nikolaev Viacheslav, M. Katschinski Dörthe, Redox Imaging Using Cardiac Myocyte-Specific Transgenic Biosensor Mice. Circulation Research 119, 1004–1016 (2016).

53. K. Kojer, J. Riemer, Balancing oxidative protein folding: The influences of reducing pathways on disulfide bond formation. Biochimica et Biophysica Acta (BBA) - Proteins and Proteomics 1844, 1383–1390 (2014).

54. J. Riemer, N. Bulleid, J. M. Herrmann, Disulfide formation in the ER and mitochondria: two solutions to a common process. Science 324, 1284–1287 (2009).

55. M. J. Saaranen, L. W. Ruddock, Disulfide Bond Formation in the Cytoplasm. Antioxidants & Redox Signaling 19, 46–53 (2012).

56. H. S. Chung, S. B. Wang, V. Venkatraman, C. I. Murray, J. E. Van Eyk, Cysteine oxidative posttranslational modifications: emerging regulation in the cardiovascular system. Circ Res 112, 382–392 (2013).

57. F. Cuello, I. Wittig, K. Lorenz, P. Eaton, Oxidation of cardiac myofilament proteins: Priming for dysfunction? Molecular Aspects of Medicine 63, 47–58 (2018).

58. C. X. C. Santos, N. Anilkumar, M. Zhang, A. C. Brewer, A. M. Shah, Redox signaling in cardiac myocytes. Free Radical Biology and Medicine 50, 777–793 (2011).

59. J. A. Rivas-Pardo, E. C. Eckels, I. Popa, P. Kosuri, W. A. Linke, J. M. Fernandez, Work Done by Titin Protein Folding Assists Muscle Contraction. Cell reports 14, 1339–1347 (2016).

60. A. Brynnel, Y. Hernandez, B. Kiss, J. Lindqvist, M. Adler, J. Kolb, R. van der Pijl, J. Gohlke, J. Strom, J. Smith, C. Ottenheijm, H. L. Granzier, Downsizing the molecular spring of the giant protein titin reveals that skeletal muscle titin determines passive stiffness and drives longitudinal hypertrophy. eLife 7, e40532 (2018).

61. W. A. Linke, M. R. Stockmeier, M. Ivemeyer, H. Hosser, P. Mundel, Characterizing titin’s I-band Ig domain region as an entropic spring. Journal of Cell Science 111, 1567–1574 (1998).

62. E. C. Eckels, S. Haldar, R. Tapia-Rojo, J. A. Rivas-Pardo, J. M. Fernandez, The Mechanical Power of Titin Folding. Cell reports 27, 1836–1847 e1834 (2019).

63. Y.-Q. Zhou, F. S. Foster, R. Parkes, S. L. Adamson, Developmental changes in left and right ventricular diastolic filling patterns in mice. American Journal of Physiology-Heart and Circulatory Physiology 285, H1563–H1575 (2003).

64. S. Kötter, M. Kazmierowska, C. Andresen, K. Bottermann, M. Grandoch, S. Gorressen, A. Heinen, J. Moll, J. Scheller, A. Gödecke, J. W. Fischer, J. P. Schmitt, M. Krüger, Titin-Based Cardiac Myocyte Stiffening Contributes to Early Adaptive Ventricular Remodeling After Myocardial Infarction. Circulation Research, 1017–1029 (2016).

65. C. Suay-Corredera, M. R. Pricolo, E. Herrero-Galan, D. Velazquez-Carreras, D. Sanchez-Ortiz, Garcia-Giustiniani, J. Delgado, J. J. Galano-Frutos, H. Garcia-Cebollada, S. Vilches, F. Dominguez, M. Sabater Molina, R. Barriales-Villa, G. Frisso, J. Sancho, L. Serrano, P. Garcia-Pavia, L. Monserrat, J. Alegre-Cebollada, Protein haploinsufficiency drivers identify MYBPC3 mutations that cause hypertrophic cardiomyopathy. medRxiv, 2020.2005.2004.20087726 (2020).

66. C. Chauveau, J. Rowell, A. Ferreiro, A Rising Titan: TTN Review and Mutation Update. Human mutation 35, 1046–1059 (2014).

67. K. M. Meurs, S. G. Friedenberg, J. Kolb, C. Saripalli, P. Tonino, K. Woodruff, N. J. Olby, B. W. Keene, D. B. Adin, O. L. Yost, T. C. DeFrancesco, S. Lahmers, S. Tou, G. D. Shelton, H. Granzier, A missense variant in the titin gene in Doberman pinscher dogs with familial dilated cardiomyopathy and sudden cardiac death. Human Genetics 138, 515–524 (2019).

68. L. Begay Rene, S. Graw, G. Sinagra, M. Merlo, D. Slavov, K. Gowan, L. Jones Kenneth, G. Barbati, A. Spezzacatene, F. Brun, A. Di Lenarda, E. Smith John, L. Granzier Henk, L. Mestroni, M. Taylor, n. null, Role of Titin Missense Variants in Dilated Cardiomyopathy. Journal of the American Heart Association 4, e002645 (2015).

69. Y. M. Pinto, P. M. Elliott, E. Arbustini, Y. Adler, A. Anastasakis, M. Böhm, D. Duboc, J. Gimeno, P. de Groote, M. Imazio, S. Heymans, K. Klingel, M. Komajda, G. Limongelli, A. Linhart, J. Mogensen, J. Moon, P. G. Pieper, P. M. Seferovic, S. Schueler, J. L. Zamorano, A. L. P. Caforio, P. Charron, Proposal for a revised definition of dilated cardiomyopathy, hypokinetic non-dilated cardiomyopathy, and its implications for clinical practice: a position statement of the ESC working group on myocardial and pericardial diseases. European Heart Journal 37, 1850–1858 (2016).

70. L. Padrón-Barthe, M. Villalba-Orero, M. Gómez-Salinero Jesús, F. Domínguez, M. Román, J. Larrasa-Alonso, P. Ortiz-Sánchez, F. Martínez, M. López-Olañeta, E. Bonzón-Kulichenko, J. Vázquez, C. Martí-Gómez, J. Santiago Demetrio, B. Prados, G. Giovinazzo, V. Gómez-Gaviro María, S. Priori, P. Garcia-Pavia, E. Lara-Pezzi, Severe Cardiac Dysfunction and Death Caused by Arrhythmogenic Right Ventricular Cardiomyopathy Type 5 Are Improved by Inhibition of Glycogen Synthase Kinase-3β. Circulation 140, 1188–1204 (2019).

71. J. Herron Todd, D. Rocha Andre Monteiro, F. Campbell Katherine, D. Ponce-Balbuena, B. C. Willis, G. Guerrero-Serna, Q. Liu, M. Klos, H. Musa, M. Zarzoso, A. Bizy, J. Furness, J. Anumonwo, S. Mironov, J. Jalife, Extracellular Matrix–Mediated Maturation of Human Pluripotent Stem Cell–Derived Cardiac Monolayer Structure and Electrophysiological Function. Circulation: Arrhythmia and Electrophysiology 9, e003638 (2016).

72. L. Sala, J. van Meer Berend, G. J. Tertoolen Leon, J. Bakkers, M. Bellin, P. Davis Richard, C. Denning, A. E. Dieben Michel, T. Eschenhagen, E. Giacomelli, C. Grandela, A. Hansen, R. Holman Eduard, R. M. Jongbloed Monique, M. Kamel Sarah, D. Koopman Charlotte, Q. Lachaud, I. Mannhardt, P. H. Mol Mervyn, D. Mosqueira, V. Orlova Valeria, R. Passier, C. Ribeiro Marcelo, U. Saleem, L. Smith Godfrey, L. Burton Francis, L. Mummery Christine, MUSCLEMOTION. Circulation Research 122, e5–e16 (2018).

73. C. Pimenta-Lopes, C. Suay-Corredera, D. Velázquez-Carreras, D. Sánchez-Ortiz, J. Alegre-Cebollada, Concurrent atomic force spectroscopy. Communications Physics 2, 91 (2019).

74. E. Gasteiger, C. Hoogland, A. Gattiker, S. e. Duvaud, M. R. Wilkins, R. D. Appel, A. Bairoch, in The Proteomics Protocols Handbook, J. M. Walker, Ed. (Humana Press, Totowa, NJ, 2005), pp. 571–607.

