## Supplementary Material for "Conserved cysteines in titin sustain the mechanical function of cardiomyocytes"

- **Supplementary File S1 (separate file)**
- **Supplementary Figures S1-S5 (this document)**
- **Supplementary Notes S1-S4 (this document)**
- **Supplementary Tables S1-S5 (this document)**

¶ EHG and FD contributed equally and are joint first authors.

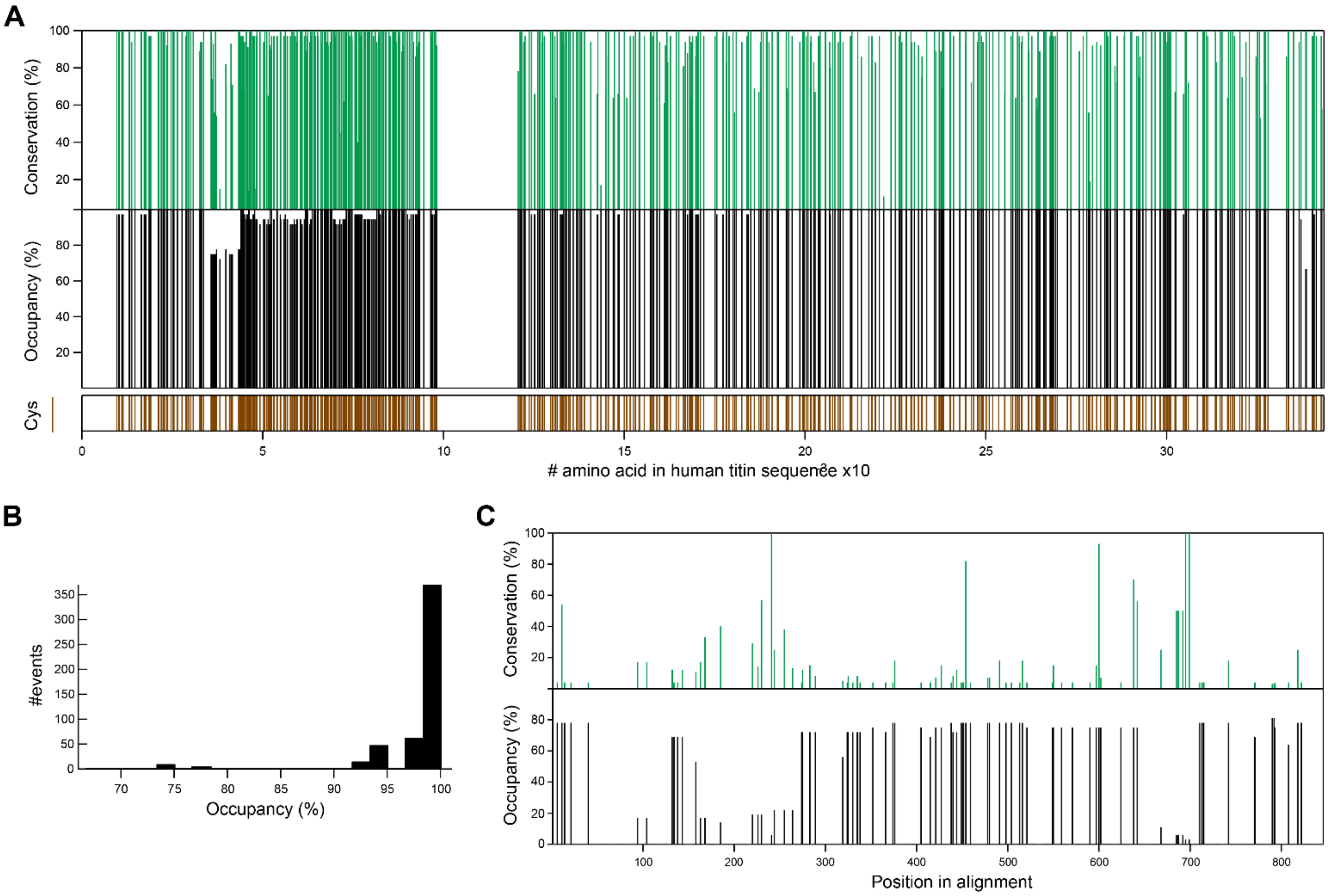


**Supplementary Figure S1. Evolutionary conservation of titin cysteines. A:** Percentage of conservation (top, green) and occupancy (middle, black) of every cysteine in human titin (bottom, brown, Uniprot Q8WZ42-1) in a sequence alignment including titins from 36 species (**Supplementary File S1**). **B:** Distribution of occupancy of all cysteines in human titin. **C:** Percentage of conservation (top, green) and occupancy (bottom, black) of every N2Bus cysteine in the interspecies alignment.

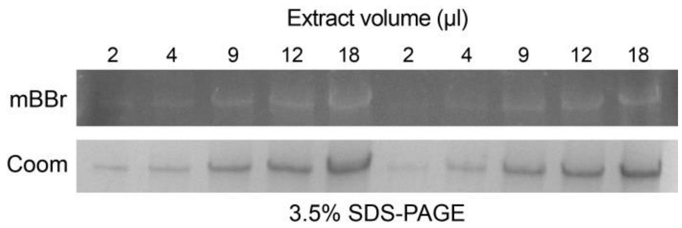


**Supplementary Figure S2. Linearity of mBBr cysteine oxidation signal**. mBBr and Coomassie (Coom) stainings of titin bands resulting from loading different volumes of the same mouse heart extract in a 3.5% SDS-PAGE gel. Quantification is shown in Figure 2B-D.

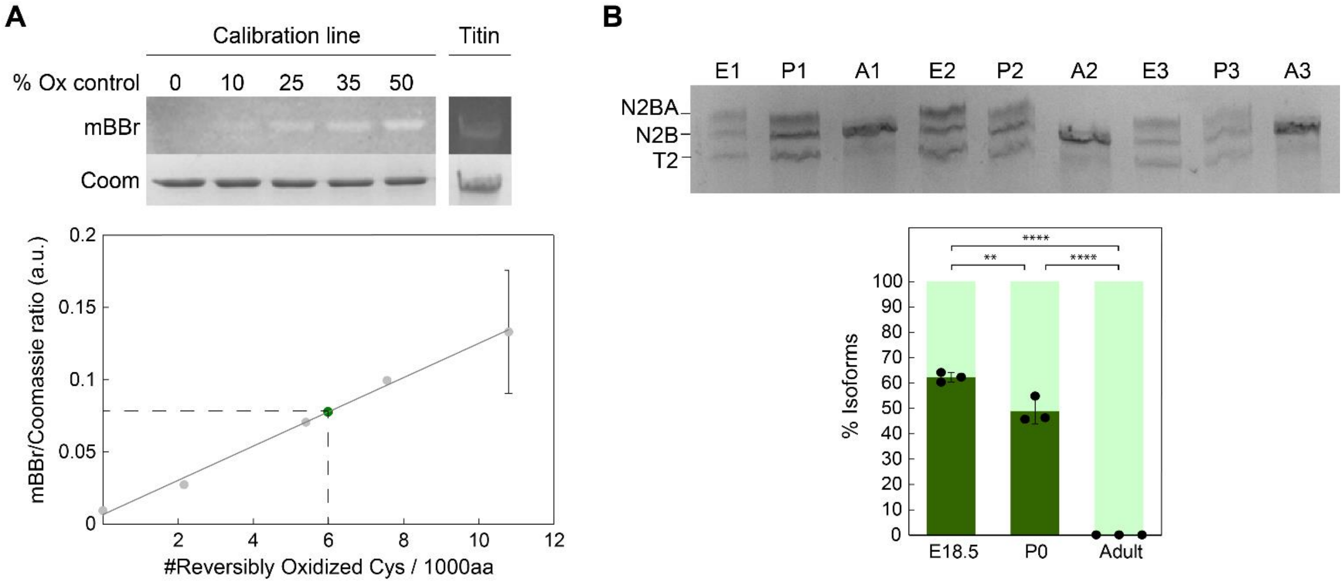


**Supplementary Figure S3. Redox calibration curve and quantification of titin isoforms in perinatal and adult mouse samples. A:** Samples containing known fractions of oxidized (I91–32/75)$_8$ polyprotein are used to build a calibration line of Coomassie-normalized mBBr fluorescence versus number of reversibly oxidized cysteines per 1000 amino acids (12% SDS-PAGE gel). The corresponding signals coming from the titin bands of interest (3.5% SDS-PAGE gel) are interpolated in the curve (green dot), which allows averaging results from independent experiments. **B**: Quantification of the proportion of N2BA (dark green) and N2B (light green) titin isoforms in three different embryo (18.5 days, E), P0 (P) and adult (A) mice using a hybrid 2% acrylamide/1.5% agarose SDS-PAGE/agarose gel, which was run at 4ºC and 4-8 mA for 3.5 hours. T2 is a typical degradation product of titin (*75*). p(E18.5 *vs* P0) = 0.005, p(E18.5 *vs* Adult) < 0.0001, p(P0 *vs* Adult) < 0.0001 (ANOVA and Tukey's multiple comparisons test). Errors are SD.

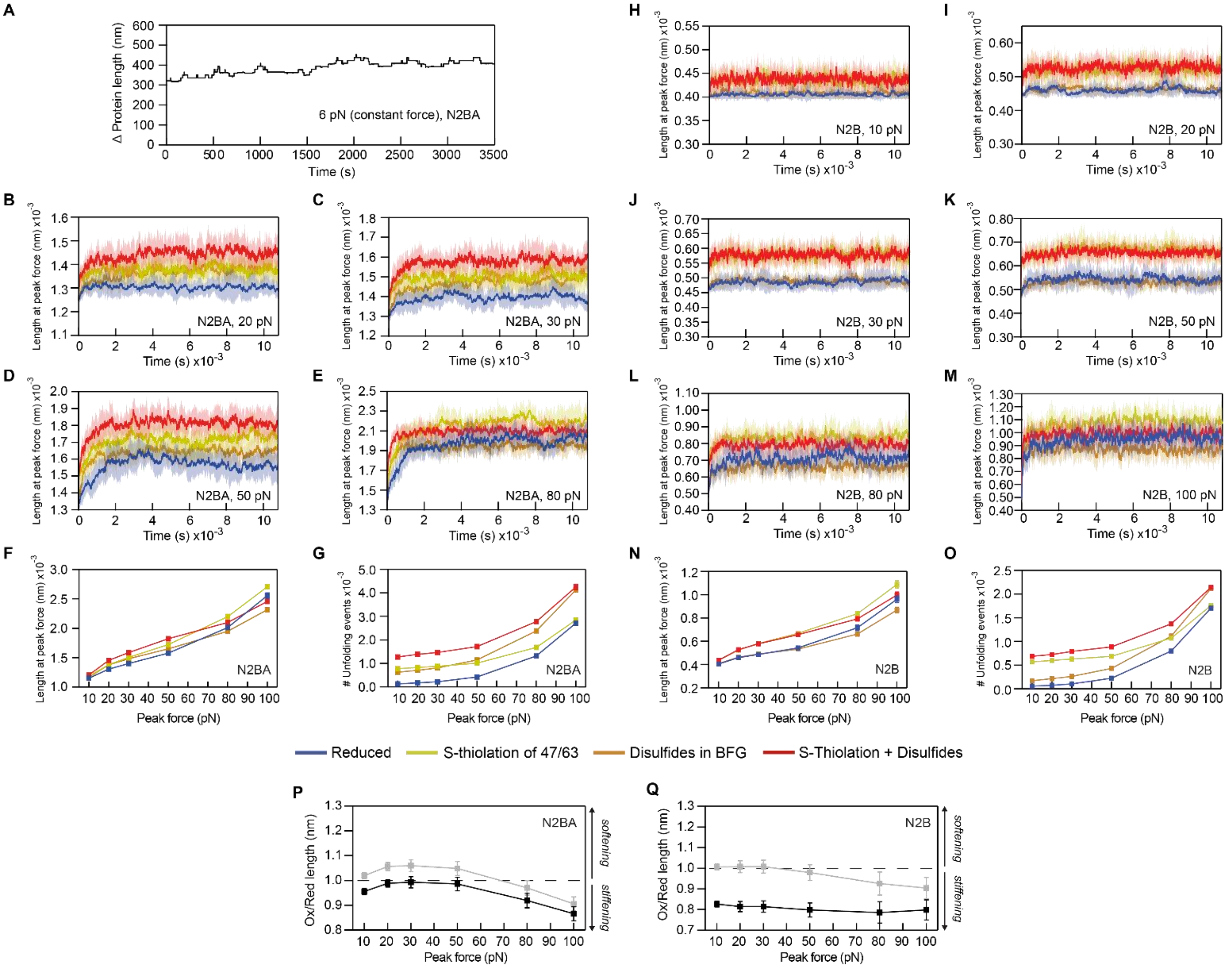


**Supplementary Figure S4. Monte Carlo simulations. A**: Monte Carlo simulations in which a virtual I-band titin is pulled at a constant force of 6 pN. Results are qualitatively similar to experimental observations (*33*). **B-E**: Length of N2BA titin at peak force during simulations in which titin is reduced, or oxidized by disulfides, S-thiolation adducts or both (color code at the bottom of the figure; peak forces are shown in the insets). For subsequent analyses, we considered times longer than 1 hour, when the length of titin fluctuates around steady state values. **F**: Length of N2BA titin at different peak forces. **G**: Cumulative number of unfolding events in simulations of N2BA titin at different peak forces. **H-M**: Length of N2B titin at peak force during simulations in which titin is reduced, or oxidized by disulfides, S-thiolation adducts or both (color code at the bottom of the figure). For subsequent analyses, we considered times longer than 1 hour. **N**: Length of N2B titin at different peak forces. **O**: Cumulative number of unfolding events in simulations of N2B titin at different peak forces. **P**: Ratio of oxidized *vs* reduced N2BA titin length at different peak forces considering (black) or not (grey, same data as in **Figure 6E**) that the N2Bus region contains disulfides. **Q**: Ratio of oxidized *vs* reduced N2B titin length at different peak forces considering (black) or not (grey, same data as in **Figure 6G**) that the N2Bus region contains disulfides. In panels B-G, n = 10 simulations, and error bars or shaded areas represent SD.

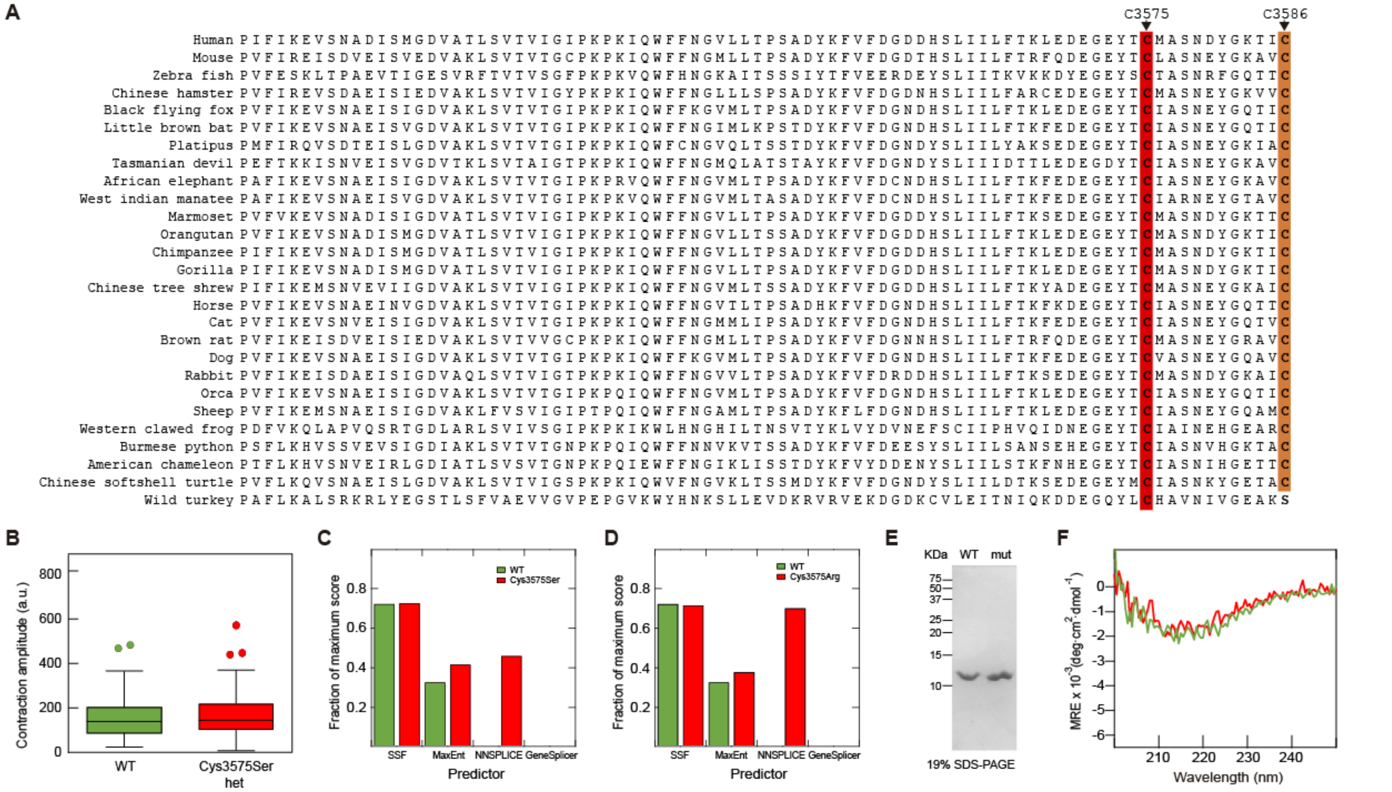


**Supplementary Figure S5. Characterization of DCM mutants targeting Cys3575 of titin. A**: Interspecies sequence alignment of human I21 titin domain. Cysteines 3575 and 3586 are highlighted in red and orange, respectively. **B**: Box-and-whiskers plots showing amplitude of contraction of WT (green, n = 68) and heterozygous Cys3575Ser (red, n = 78) hiPSC-induced cardiomyocytes (p=0.5326, Mann-Whitney). Box encompasses data between quartiles 1 and 3 and horizontal lines represent the median of the distributions. Whiskers and outliers were calculated using Tukey's method as implemented in Graph Pad. **C**, **D**: Maximum splicing scores from different predictors implemented in Alamut Visual (Interactive Biosoftware) for WT (green) and variants affecting Cys3575 (red). Only NNSplice identifies the possibility of a new splicing site induced by the mutations with a high score. This limited degree of prediction is typical of false positives that results in no experimental alteration of splicing (*65*). **E**: 19% SDS-PAGE of purified recombinant WT and Cys3575Ser (mut) I21 domains. **F**: Far-UV circular dichroism spectra at 85ºC of WT (green) and Cys3575Ser (red) recombinant I21 protein domains.

### Supplementary Note S1. Titin model used for Monte Carlo simulations

Titin length as a function of end-to-end force was modeled on the basis of the Freely Jointed Chain (FJC) model of polymer elasticity (*76*), which was applied independently to Ig domains and the random coil regions of the protein. The total length of titin was then calculated by adding up the contribution of the different protein regions. The Freely Jointed Chain model calculates molecular extension as a result of end-to-end force following

$$Length_{FJC}(F) = L_C\left[coth\left(\frac{F \cdot L_k}{k_B \cdot T}\right) - \frac{k_B \cdot T}{F \cdot L_k}\right]$$

where $F$ is the applied force, $L_c$ is the contour length, $L_k$ is the Kuhn length, $T$ is the absolute temperature and $k_B$ is the Boltzmann constant.

All folding and unfolding events were considered to depend on the applied force according to Bell's model (*77*). Disulfide isomerization reactions were modelled in a similar way, but with the additional restriction that they can only take place in the unfolded state (*21*). The probability of unfolding, refolding or isomerization events was calculated according to

$$P_{a \to b} = N_a * dt * \alpha_0 * e^{\frac{F\,\Delta x}{k_B T}}$$

where $a$ and $b$ are 2 possible states of the domain, $N_a$ is the number of domains in the state $a$, $dt$ is the simulation step, $\alpha_0$ is the transition rate at zero force and $\Delta x$ is the distance to the transition state. Definition of titin regions and parameters used in the simulations can be found in **Supplementary Tables S1-S4**. Transition rates are in the range of those previously determined for specific segments of titin (*19*, *21*, *22*), but were further adjusted so that simulations reproduce the folding/unfolding dynamics of native Ig domains stretched at low forces (*33*). We also considered that modifications affect the maximum folding capacity of the domains (*22*) (**Supplementary Table S5**).

**Supplementary Note S2. Recombinant I21 WT and Cys3575Ser *E.coli*-codon-optimized cDNA and protein sequences.** The codon codifying for Cys3575 is shown in red; in the mutation, the first T is substituted for an A. Residues corresponding to titin (amino acid positions 3500 to 3598 according to Uniprot Q8WZ42) are indicated in green and extra residues added in the cloning appear in black.

**cDNA:**

ATGAGAGGATCGCATCACCATCACCATCACGGATCCGGCACAGGCCCGATCTTTATCAAAGAAGTTAG
CAACGCAGATATTAGCATGGGTGATGTTGCAACCCTGAGCGTTACCGTTATTGGTATTCCGAAACCGAA
AATCCAGTGGTTTTTTAACGGTGTTCTGCTGACCCCGAGCGCAGATTACAAATTCGTTTTTGATGGTGAT
GATCACAGCCTGATTATCCTGTTTACCAAACTGGAAGATGAAGGCGAATATACC**TGT**ATGGCAAGCAA
TGATTATGGCAAAACCATTTGTAGCGCCTACCTGAAAATCAATAGTAAGGGCGAAGGTAGATCTTAA

**Protein:**

MRGSHHHHHHGSGTGPIFIKEVSNADISMGDVATLSVTVIGIPKPKIQWFFNGVLLTPSADYKFVFDGDDHSL
IILFTKLEDEGEYT**C**MASNDYGKTICSAYLKINSKGEGRS-

**Supplementary Note S3. Sequence identification codes for the analysis of interspecies conservation of titin residues**

| | |
|---|---|
| A5X6X5 | *Danio rerio (Zebrafish)* |
| XP_025759277 | *Oreochromis niloticus (Tilapia)* |
| XP_031748776 | *Xenopus tropicalis (Western clawed frog)* |
| XP_025018685 | *Python bivittatus (Burmese python)* |
| XP_016851936 | *Anolis carolinensis (American chameleon)* |
| XP_025038559 | *Pelodiscus sinensis (Chinese softshell turtle)* |
| XP_019335759 | *Alligator mississippiensis (American alligator)* |
| M7AMC7 | *Chelonia mydas (Green sea turtle)* |
| XP_032605298 | *Taeniopygia guttata (Zebrafinch)* |
| XP_027317399 | *Anas platyrhynchos (Wild duck)* |
| XP_031410147 | *Meleagris gallopavo (Wild turkey)* |
| XP_027642041 | *Falco peregrinus (Peregrine falcon)* |
| XP_016155182 | *Ficedula albicollis (Collared flycatcher)* |
| XP_028928092 | *Ornithorhynchus anatinus (Platypus)* |
| XP_031816250 | *Sarcophilus harrisii (Tasmanian devil)* |
| XP_023398816 | *Loxodonta africana (African elephant)* |
| XP_004375522 | *Trichechus manatus (West Indian manatee)* |
| L5K2L4 | *Pteropus alecto (Black flying fox)* |
| G1P5X9 | *Myotis lucifugus (Little brown bat)* |
| XP_023115277 | *Felis catus (Cat)* |
| XP_022270508 | *Canis lupus (Dog)* |
| XP_019654795 | *Ailuropoda melanoleuca (Giant panda)* |
| A0A5F5PSR4 | *Equus caballus (Horse)* |
| XP_004267426 | *Orcinus orca (Orca)* |
| DAA32835 | *Bos taurus (Cattle)* |
| XP_027820594 | *Ovis aries (Sheep)* |
| XP_017198704 | *Oryctolagus cuniculus (Rabbit)* |
| G3HAC6 | *Cricetulus griseus (Chinese hamster)* |
| A2ASS6 | *Mus musculus (Mouse)* |
| XP_017456475 | *Rattus norvegicus (Brown Rat)* |
| XP_027628916 | *Tupaia chinensis (Chinese tree shrew)* |
| A0A5F4VS40 | *Callithrix jacchus (Marmoset)* |
| A0A2J8VRG3 | *Pongo abelii (Orangutan)* |
| XP_030864472 | *Gorilla gorilla (Gorilla)* |
| A0A2J8PRF8 | *Pan troglodytes (Chimpanzee)* |
| Q8WZ42 | *Homo sapiens (Human)* |

**Supplementary Note S4. Alignment of the 144 murine titin Ig domains in the canonical sequence Uniprot A2ASS6.** The position of the first amino acid of each domain is indicated at the beginning of each row and cysteines are highlighted in yellow. The position of the structurally conserved cysteines is indicated on top of the alignment according to the nomenclature in the main text. An equivalent alignment for human titin has been published (*21*).

```
                                                   CysB                                                         Cys47                Cys63     CysF          CysG
00006 ----------------PMFT---QPLQ-S-VVVLEGS----TATFE-AH--VSGSPV-PEVSWFRDGQ--V-IS-TSTLP---------------------------GVQISF---S--DGRARLMIPAVTKANSGRYSLRATNGSGQA-TSTAELLVT--
00104 ----------------PNFS---QRLQ-S-MTVRQGS----QVRLQ-VR--VTGIPT-PVVKFYRDGA--E-IQ-SS--L---------------------------DFQISQ---E--GDLYSLLIAEAYPEDSGTYSVNATNSVGRA-TSTAELV----
00946 ----------------PTLV---SGLK-N-VTVIEGE----SVTLE-CH--ISGYPS-PKVTWYREDY--Q-IE-SS--I---------------------------DFQITF---Q--GGIARLMIREAFAEDSGRFTCSAVNEAGTV-STSCYLA----
01084 ----------------PFFI---SKPV-V-QKLVEGG----SVVFE-CQ--IGGNPK-PHVYWKKSGV--P-LT-TG----------------------Y-------RYKVSY-NK-Q-TGECRLVISMTFADDAGEYTIVIRNKHGET-SASASLL----
01293 ----------------SGFD---IRIK-N-YRILEGM----GVTFH-CK--MSGYPL-PKIAWYKDGK--R-IR-HG--E---------------------------RYQMDF---LQ-DGRASLRIPVVLPEDEGIYTAFASNIKGNA-ICSGKLYVE--
01463 ----------------PVFV---LKPA-S-FKCLEGQ----TARFD-LK--VVGRPM-PETFWFHNGQ--Q-IV-ND--Y---------------------------T--HKVVIK-E-DGTQSLIIVPASPSDSGEWTVVAQNRAGKS-TISVTLT----
01562 ----------------PAFVEKL-----KNVNIKEGS----RLEMK-VR--ATGNPN-PDIVWLKNSD--I-IV--P--H-------------------K-----YPR--IRI-E-GT-RGEAALKIDSIISQDSAWYTATAINKAGRD-TTRCKVN----
01709 ----------------PFFK---KKLT-S-LRLKRFG----PAHFE-CRLTPIGDPT-MVVEWLHDGK--P-LE-AA--N---------------------------R--LRM-IN-E-FGYCSLDYGAAYSRDSGVITCRATNKYGTD-HTSATLI----
01847 ----------------PDIV---LFPE-P-VRVLEGE----TARFR-CR--VTGYPQ-PKVNWYLNGQ--L-IR-KS--K---------------------------R--F--RVR-Y-DGIHYLDIVDCKSYDTGEVKVTAENPEGVT-EHKVKLE----
02084 ----------------PKIF---ERIQ-S-QTVGQGS----DAHFR-VR--VVGKPD-PECEWYKNGV--K-IE-RS--D---------------------------R--IYWYWP-E-DNVCELVIRDVTAEDSASIMVKAINIAGET-SSHAFLL----
02177 -------------KQLITFT---QELQ-D-VVAKEKD---TMATFE-CE--TSEPFI--KVKWYKDGI--E-VH-AG--D---------------------------K--YRM-H-SD-RKVHFLSVLTIDTSDAEDYSCVLVEDENIK--TTAKLIV---
02270 -------------GAVVEFV---KELQ-D-IEVPESY----SGELE-CI--ISPENI--EGKWYHNDV--E-LK-SN--G---------------------------K--YSI-T-SR-RGRQNLTVKDVTKEDQGEYSFVVDGKK-----TTCKLK----
02359 -------------PRPIAIL---QGLS-D-QKVCEGD----IVQLE-VK--VSLENV--EGVWMKDGQ--E-VQ-HS--D---------------------------R--VHI-V-ID-KQSHMLLIEDMTKEDAGNYSFTIPALGLST-SGNVSVY----
02436 -PALGLSTSGNVSVYSVDVI---TPLK-D-VNVIEGT----KAVLE-CK--VSVPDV-TSVKWYLNDE--Q-IK-PD--D---------------------------R--VQS-I-VK-GTKQRLVINRTHASDEGPYKLMVGRVE-----TSCNLS----
02626 ----------------GAIS---TPLT-D-QTVAESQ----EAVFE-CE--VANPES--EGEWLKDGK--H-LA-LS--N---------------------------N--FRG-E-SD-GHKRRLVIAAAKLDDAGEYTYKVATSK-----TSAKLK----
02886 --------------ETLHIT---KTMK-S-IEVPETK----AASFE-CE--VSHFNV--PSMWLKNGV--E-IE-MS--E---------------------------K--FKI-V-VQ-GKLHQLIIMNTSTEDSAEYTFVCGNDQ-----VSATLT----
03064 ----------------IEFR---KHIK-D-IKVLEKK----RAMFE-CE--VSEPDI--TVQWMKDGQ--E-LQ-IA--D---------------------------R--IKI-Q-KE-KYVHRLLIPSTRMSDAGKYTVVAGGNMSTA--NLF-------
03245 ----------------PQVL---QELQ-P-VTVQSGK----PARFC-AV--ISGRPQ-PKISWYKEEQ--L-LS-TG--F---------------------------K--CKF-LH-D-GQEYTLLLIEAFPEDAAVYTCEAKNDYGVA-TTSASLS----
03350 ---------------PPAIV---TPLQ-D-TVTSEGR----PARFQ-CQ--VSGTDL--KVSWYCKDK--K-IK-PS--R---------------------------F--FRM-TQ-F-EDTYQLEIAEAYPEDEGTYAFVANNAVGQV-SSTATLR----
03509 ----------------PVFI---REIS-D-VEISVED----VAKLS-VT--VTGCPK-PKIQWFFNGM--L-LT-PS--A---------------------------DYKFVF---D--GDTHSLIILFTRFQDEGEYTCLASNEYGKA-VCSAHLRIS--
03625 ----------------PYFF---KELK-P-VHCGPGI----PAVFE-YS--VHGEPA-PTVLWFKEDM--P-LY-TS--V---------------------------C--YTI-IH-SPDGSGTFIVNDPQRGDSGLYLCKAQNLWGES-TCAAELLVL--
04251 ----------------PAVL---KHLV-D-TISEEGD----TVHLT-SS--ISNAK---EVHWYFKGN--L-VP-SD--G---------------------------K--FKC-LK-E-QNAYTLVIEAVKTEDEGEYVCEASNDSGKA-KTSAKLT----
04344 ----------------PVIK---RRIE-P-LEVALGH----LAKFT-CE--IQGAPN-VRFQWFKAGR--E-IY-ES--D---------------------------KCSIRS---S--NYVSSLEILRTQVVDCGEYTCKASNEYGSV-SCTATLT----
04439 ----------------PTFL---SRPK-A-LTTFVGK----AAKFL-CT--VSGTPV-IEIIWQKDGA--A-LS-PS--P---------------------------DCRVTD---A--DNKHSLELSNLTVQDRGIYSCKASNKFGAD-ICQAELT----
04532 ----------------PHFI---KELE-A-VQSAINK----KIHLE-CQ--VDEDRK-VTITWSKDGQ--K-LP-AG--K---------------------------DYKIYF---E--DKIASLEIPLAKLKDSGTYTCTASNEAGSS-SSSAAVA----
04625 ----------------PSFV---KKVD-PSYLMLPGE----SARLH-CK--LKGSPV-IQVTWFKNNK--E-LS-ES--N---------------------------TVRMSF---V--NSEAILDITDVKVDDSGTYSCEATNDVGSD-SCSTEVVIK--
04719 ----------------PSFI---KTLE-P-ADIVRGA----NALLQ-CE--IAGTGP-FEVNWFKDKK--Q-IR-SS--K---------------------------KYRLFT---Q--KTFVYLEISSFNSADVGDYECVVANEVGKC-GCVATHL----
04812 ----------------PTFV---KKVD-D-FTALAGQ----TVTLQ-AA--VRGSEP-ISVMWMKGQE--V-IK-ED--G---------------------------KIKMSF---S--NGVAVLTIPDVQISLGGKYTCLAENEAGSQ-TSVG-------
04904 ---------------PAKII---ERAE-L-IQVTAGD----PATLE-YT--VSGTPE-LKPKWYKDGR--P-LV-AS--K---------------------------KYRISF---K--NNIAQLKFYSAELHDSGQYTFEISNEVGSS-SCETTFT----
05001 ----------------PLFT---KPLR-N-VDSVVGG----ACRLD-CK--IAGSLP-MRVSWFKDGK--E-LT-AS--D---------------------------RYQIAF---V--EGTASLEISRVDMNDAGNFTCRATNSVGSK-DSSGALI----
05094 ----------------PSFV---TKPG-S-RDVLPGS----AVCLK-SA--FQGSAP-LTIKWFKGDK--E-LV-SG--G---------------------------SCYITK---E--TSESSLELYAVKTSDSGTYTCKVSNVAGSV-ECSADLF----
05186 ---------------PATFI---EKLE-PSQLLKKGD----GTQLA-CK--VTGTPP-IKITWFANDR--E-LR-ES--S---------------------------KHKMSF---A--ESTAVLRLTDVAIEDSGEYMCEAQNEAGSD-HCTGIVI----
05281 ----------------PYFT---KEFK-S-IEVLKEY----DVMLL-AE--VAGTPP-FEITWFKDNT--T-LR-SG--R---------------------------KYKTFL---Q--DQLVSLQVLKFVAADAGEYQCRVTNEVGSS-TCSARVT----
05374 ----------------PSFI---KKIE-A-TSSLRGG----TAAFQ-AT--LKGSLP-ITVTWLKDND--E-IT-ED--D---------------------------NIRMTF---E--NNVASLYLSGIEVKHDGKYVCQAKNDAGIQ-RCSALLS----
05466 ---------------PATIM---EEAV-S-IDVTQGD----PATLQ-VK--FSGTKE-ISAKWFKDGQ--E-LT-LG--P---------------------------KYKISV---T--DTVSILKIISTEKKDSGEYTFEVQNDVGRS-SCKASIN----
05563 ----------------PSFT---KKLR-K-MDSIKGS----FIDLE-CI--VAGSHP-ISIQWFKDDQ--E-IS-AS--D---------------------------KHKFSF---H--DNTAFLEISQLEGTDSGTYTCSATNKAGHS-QCSGHLT----
05656 ----------------PYFV---EKPQ-S-QDVNPGT----RVQLK-AL--VGGTAP-MTIKWFKDNK--E-LH-PG--A---------------------------ARSVWK---D--DTSTILELFSAKAADSGTYICQLSNDVGTT-SSKATIF----
05749 ----------------PQFI---KKPS-PVLVLRNGQ----STTFE-CQ--VTGTPE-IRVSWYLDGN--E-IT-DL--R---------------------------KYGISF---V--DGLATFQISNARVENSGTYVCEARNDAGTA-SCSIELK----
05842 ---------------PPIFI---RELE-P-VEVVKDS----DVELE-CE--VMGTTP-FEVTWLKNNK--E-IR-SG--K---------------------------KYTMSE---K--MSVFYLHITKCDPSDVGEYQCIIANEGGSC-ACSARVA----
05936 ----------------PSFI---KKIE-N-VTTVLKS----SATFQ-ST--VAGSPP-ISITWLKDDQ--I-LE-EN--D---------------------------NVHISF---E--DSVATLQVRSVDNGHSGRYTCQAKNESGIE-RCYAFLL----
06028 ---------------PAQII---EKAK-S-VDVTEKD----PVTLE-CV--VAGTPE-LKVKWLKDGK--Q-IV-PS--R---------------------------YFSMSF---E--NNVASFRIQSVMKQDSGQYTFKVENDFGSS-SCDAYLR----
06125 ----------------PSFT---KKLT-K-MDKVLGS----SIHME-CK--VSGSLP-ISAQWFKDGK--E-IS-TS--A---------------------------KYRLVC---H--ENTVSLEVSNLELEDTANYTCKVSNVAGDN-ACSGILT----
06218 ----------------PSFL---VKPE-R-QQAIPDS----TVEFK-AV--LKGTPP-FKIKWLKDDV--E-LV-SG--P---------------------------KCFIGL---E--GSTSFLNLYSVDSSKTGQYTCQVTNDVGSD-SCTTMLLVT--
06311 ----------------PKFV---KKLE-ASKIIKAGD----SARLE-CK--ITGSPE-IQVVWYRNEH--E-LT-AS--D---------------------------KYQMTF---I--DSVAVIQMNSLGTEDSGDFICEAQNPAGST-SCSTKVIV---
06405 ----------------PVFS---SFPP-I-VETLKNT----EVSLE-CE--LSGTPP-FEVVWYKDKR--Q-LR-SS--K---------------------------KYKVAS---K--NFHASIHILNVESTDIGEYHCKAQNEVGSD-ACVCAVK----
06498 ----------------PKFI---SKLN-S-LTVVAGE----PAELQ-AS--IEGAQP-ISVQWLKEKE--EVIR-ES--E---------------------------NIRISF---V--NNVATLQFAKVEPANAGKYICQVKNDGGVR-ENMATLT----
06591 ---------------PAVII---EKAG-S-MTVTVGE----TCALE-CK--VAGTPE-LSVEWYKDGK--L-LT-SS--Q---------------------------KHKFSF---Y--NKISSLKILSVEKEDAGTYTFQVQNNVGKS-SCTAVVDVS--
06688 ----------------PSFT---RRLK-D-TGGVLGT----SCILE-CK--VAGSSP-ISIAWFHEKT--K-IV-SG--A---------------------------KYQTTF---S--DNVCTLQLNSLDSSDMGSYTCVAANVAGSD-ECRALLT----
06781 ----------------PSFV---KEPE-P-LEVLPGK----NITFT-SV--IRGTPP-FKVGWFRGAR--E-LV-KG--N---------------------------RCNIYF---E--DTVAELELFNIDISQSGEYTCVVSNNAGQA-SCTTRLF----
06873 ---------------PATFV---KKLS-D-HSVEPGK----SIILE-GT--YTGTLP-ISVTWKKDGV--S-IT-PS--E---------------------------RCNIVT---T--EKTCILEILSSTKGDAGHYSCEIENEAGRD-ACDALVS----
06966 ---------------PPYFV---TELE-P-LEASVGD----SVSLQ-CQ--VAGTPE-ITVSWFKGDT--K-LR-ST--P---------------------------EYRTYF---T--NNVATLVFNKVSINDSGEYTCMAENSIGTA-ASKTIF-----
07063 ----------------PSFA---RQLK-D-IEQTVGL----PVTLT-CR--LNGSAP-IQVCWYRDGV--L-LR-DD--E---------------------------NLQMSF---V--DNVATLKILQTDLSHSGQYSCSASNPLGTA-SSTARLT----
07159 ----------------PFFD---IKPV-S-IDVIAGE----SADFE-CH--VTGAQP-MRVTWSKDNK--E-IR-PG--G---------------------------NYTITC---V--GNTPHLRILKVGKGDSGQYTCQATNDVGKD-MCSAQLS----
07252 ----------------PKFI---KKLD-ASKVAKQGE----SIQLE-CK--ISGSPE-IKVVWFRNDS--E-LH-ES--W---------------------------KYNMSF---V--NSVALLTINEASAEDTGDYICEAHNGVGDA-SCSTALKVK--
07346 ----------------PVFT---QKPP-P-VGALKGS----DVILQ-CE--ISGTPP-FEVVWVKDRK--Q-VR-SS--K---------------------------KFKITS---K--NFDTSLHIFNLEAPDIGEYHCKATNEVGSD-TCACTVK----
07439 ----------------PRFV---KKLS-D-ASTLIGD----PVELQ-AV--VEGFQP-ISVVWLKDKG--EVIR-ES--E---------------------------NVRISF---V--DNIATLQLGSPEASQSGKYVCQIKNDAGMR-ECSAVLT----
07532 ---------------PATIV---EKPE-P-MTVTTGN----PFTLE-CV--VAGTPE-LSAKWFKDGR--E-LS-SG--S---------------------------RHHITF---V--RNLASLKIPSAEMNDKGLYTFEVENRVGKS-SCTVSVHVS--
07629 ----------------PSFV---RRLK-D-TSATLGA----SVVLE-CR--VSGSAP-ISVGWFLDGN--E-II-SS--P---------------------------KCQSSF---A--DNVCTLTLSSLEPSDTGAYTCVAANVAGQD-ESSAVLT----
07722 ----------------PSFE---QTPD-S-VEVLPGM----SLTFT-SV--IRGTPP-FKVKWFKGSR--E-LV-SG--E---------------------------ACTISL---E--DFVTELELLEVEPGQSGDYSCLVTNDAGSA-SCTTHLF----
07814 ---------------PATFV---KRLA-D-TSVETGS----PIVLE-AT--YSGTPP-ISVSWMKNEY--P-LS-QS--P---------------------------NCGITT---T--EKSSILEILESTIEDYAQYACLIENEAGQD-ICEALVS----
07907 ---------------PPYFI---EPLE-H-VEAAIGE----PITLQ-CK--VDGTPE-IRISWYKEHT--K-LR-SA--P---------------------------AYKMQF---K--NNVASLVINKVDHSDVGEYTCKAENSVGAV-ASSAVLV----
08004 ----------------PSFA---RKLK-D-VHETLGF----PVAFE-CR--INGSEP-LQVSWYKDGE--L-LK-DD--A---------------------------NLQMSF---V--HHVATLQILQTDQSHVGQYNCSASNPLGTA-SSSAKLILS--
08099 ---------------PPFFD---LKPV-S-VDLALGE----SGSFK-CH--VTGTAP-IKITWAKDNR--E-IR-PG--G---------------------------NYKMTL---V--ENTATLTVLKVAKGDAGQYTCYASNVAGKD-SCSAQLGVQ--
08193 ----------------PRFI---KKLD-QSRIVKQDE----YTRYE-CK--IGGSPE-IKVLWYKDEV--E-IQ-ES--S---------------------------KFRMSF---E--DSVAILEMHSLSVEDSGDYTCEARNAAGSA-SSSTSLK----
08287 ----------------PVFR---KKPF-P-VETLKGA----DVHLE-CE--LQGTPP-FQVSWHKDKR--E-LR-SG--K---------------------------KYKIMS---E--NLLTSIHILNVDTADIGEYQCKATNDVGSD-TCVGSVT----
08380 ----------------PQFV---KKLT-D-ISTIIGK----EVQLQ-TT--IEGAEP-ISVAWFKDKG--EIVR-ES--D---------------------------NIWISY---S--ENIATLQFSRAEPANAGKYTCQIKNDAGMQ-ECYATLS----
08473 ---------------PAAIV---EKPE-S-IKVTTGD----TCTLE-CT--VSGTPE-LSTKWFKDGK--E-LT-SD--N---------------------------KYKISF---F--NKVSGLKIINVVPGDSGVYSFEVQNPVGKD-SCKVSIQVS--
08570 ----------------PSFT---RKLK-E-TNGLSGS----SVVME-CK--VYGSPP-ISVLWFHDGN--E-IS-SG--R---------------------------KYQTTL---T--DNTCALTVNMLEEADAGDYTCIATNVAGSD-ECSAPLT----
08663 ----------------PSFV---QKPD-P-MDVLTGS----NVTFT-SI--VKGSPP-FTVSWFKGST--E-LV-PG--A---------------------------RCNVSL---Q--DSVGELELFDVDTSQSGEYTCIVSNEAGRA-SCTTRLF----
08755 ---------------PAIFV---KRLN-D-YSIEKGK----PLILE-GT--FSGTPP-ISVTWKKNGI--N-VI-AS--Q---------------------------RCNITT---T--EKSAILEILSSTVEDSGQYNCYIENASGKD-SCSAQIL----
08849 ----------------PYFV---KQLE-P-VKVTVGD----SASLQ-CQ--LAGTPE-IGVSWYKGDT--K-LR-PT--A---------------------------TCKMHF---K--NNVATLVFTQVDSSDSGEYICRAENSVGEV-SSSTFLT----
08945 ----------------PSFS---RQLR-D-VQETVGL----PVVFE-CA--VSGSEP-ISVSWYKDGK--P-LK-DS--P---------------------------NIQTSF---L--DNIATLNIFKTDRSLSGQYSCTATNPIGSA-SSGAKLILT--
09040 ---------------PPFFD---IPLA-P-MDAVVGE----SADLE-CH--VTGTQP-IKVTWAKDNR--E-IR-SG--G---------------------------NYQISY---L--ENSAHLTIVKVDKGDSGQYTCYAVNEVGKD-SCTAQLN----
09137 ----------------PSFT---KKLS-ETVEETEGN----SFKLE-GR--VAGSQP-ITVAWYKNNV--E-IH-PT--S---------------------------NCEIMF---K--NNALLLQVKRASMADAGLYTCKATNDAGSA-LCTSSIV----
09233 ---------------PPVFD---QHLA-P-VTASEGD----SVQLS-CH--VQGSEP-IRIQWLKAGR--E-VK-PS--D---------------------------RCSFSF---A--SGTAMLELKETAKADSGDYVCKASNVAGSD-TSKCKVT----
09331 -----PAAKKAAVDGKLFFV---SEPQ-S-IRVVEKT----TATFI-AK--VGGDPI-PNVKWTKGKW--R-QL-NQ--G-------------------G-------R--ILI-HQ-K-GDEAKLEIRDTTKTDSGLYRCVAFNKHGEI-ESNVNLQ----
09621 ----PHIASAKLTVIEPAWE---RHLQ-D-VTLKEGQ----TCTMT-CQ--FSVPNV--KSEWFRNGR--V-LK-PQ--G---------------------------R--VKT-E-VE-HKVHKLTIADVRAEDQGQYTCKHEDLETSA-----ELR----
09721 ---------------PIQFT---KRIQ-N-IVVSEH----QSATFE-CE--VSFDDA--IVTWYKGPT--E-LT-ES--Q---------------------------K--YNF-R-ND-GRCHYMTIHNVTPDDEGVYSVIARLEPRGEARSTAELYLT--
12903 ---------------PLKFV---KEIK-D-IVLTEAESVGSSAIFE-CL--VSPSTA--ITTWMKDGS--N-IR-ES--P---------------------------K--HRF-I-AD-GKDRKLHIIDVQLSDAGEYTCVLRLGNKEK-TSTAKLI----
13000 ---------------PVRFV---KTLEEE-VTVVKGQ----PLYLS-CE--LNKER---DVVWRKDGK--I-VVEKP--G---------------------------R--IVP-G-VI-GLMRALTINDADDTDAGTYTVTVENANNLECS----------
13361 --------------IRLKFI---SPLE-D-QTVKEGQ----TATFV-CE--LSHEKM--HVVWFKNDV--K-LH-TT--R---------------------------T--VLM-S-SE-GKTYKLEIRETTLDDISQIKAQVKNLSSTA-----NLK----
13452 ----------------PYFT---VKLH-D-KTGVEKD----EIILK-CE--VSKDV---PVKWFKDGE--E-IV-PS--P---------------------------K--HSV-K-TD-GLRRILKIKKAELKDKGEYVCDCGTDT-----TKANVT----
```

```
13628 ---------------PLIFI---TPLS-D-VKVFEKD----EAKFE-CE--VSREPK--TFRWLKGTQ--E-IT-GD--D----------------------------R--FEL-I-KD-GTRHSLVIKSAAFEDEAKYMFEAEDKR-----TSGKLI----
13807 ----------------PYFT---GKLQ-D-YTGVEKD----EVILQ-CE--ISKADA--PVKWFKDGK--E-IK-PS--K----------------------------N--VVI-K-AD-GKKRMLILKKALKSDIGQYTCDCGTDQ-----TSGKLDIEDR
13982 -------------PRVIGLL---RPLK-D-VTVTAGE----TATFD-CE--LSYEDI--PVEWYLKGK--K-LE-PN--D----------------------------K--VVT-R-SE-GRVHTLTLRDVKLEDAGEVQLTAKDFKTQA-----NLF----
14072 --------------PPVEFT---KPLE-D-QTVEEEA----TAVLE-CE--VSRENA--KVKWFKNGT--E-IL-KS--K----------------------------K--YEI-V-AD-GRVRKLIIHGCTPEDIKTYTCDAKDFK-----TSCNLN----
14161 --------------PHVEFL---RPLT-D-LQVKEKE----TARFE-CE--ISKENE--KVQWFKDGA--E-IK-KG--K----------------------------K--YDI-I-SK-GAVRILVINKCLLNDEAEYSCEVRTAR-----TSGMLT----
14250 --------------EEAVFT---KNLA-N-LEVSEGD----TIKLV-CE--VSKPGA--EVIWYKGDE--E-II-ET--G----------------------------R--FEI-L-TD-GRKRILIIQNAQLEDAGSYNCRLPSSR-----TDSKVK----
14341 ----------------AEFI---SKPQ-N-LEILEGE----KAEFV-CT--ISKESF--EVQWKRDDQ--T-LE-SG--D----------------------------K--YDI-I-AD-GKKRVLVVKDATLQDMGTYVVMVGAAR-----AAAHLT----
14427 --------------EKLRII---VPLK-D-TKVKEQQ----EVVFN-CE--VNTEGA--KAKWFRNEE--A-IF-DS--S----------------------------K--YII-L-QK-DLVYTLRIRDARLDDQANFNVSLTNHRGENVKSAANLI----
14521 --------------EDLRIV---EPLK-D-IETMEKK----SVTFW-CK--VNRLNV--TLKWTKNGE--E-VA-FD--N----------------------------R--ISY-R-ID-KYKHSLIIKDCGFPDEGEYVVTAGQDK-----SVAELLIIEA
14611 ---------------PTEFV---EHLE-D-QTVTEFD----DAVFS-CQ--LSREKA--NVKWYRNGR--E-IK-EG--K----------------------------K--YKF-E-KD-GSIHRLIIKDCRLEDECEYACGVEDRK-----SRARLF----
14789 ----------------PKIK---TADQ-D-LVVDAGQ----PLTMV-VP--YDAYPK-AEAEWFKENE--P-LS-TK--T---------------------------------V-DT-T-AEQTSFRISEAKKDDKGRYKIVLQNKHGKA-EGFINLQ----
15477 ----------------PEIFLDVKLLA-G-ITVKAGT----KIELP-AT--VTGKPE-PKITWTKADT--L-LK-PD--Q----------------------------R--ITI-E-NV-PKKSTVTITDSKRSDTGTYIIEAVNVCGRA-TAVVEVNVL--
16176 ----------------PTID---LETH-D-IVVIEGE----KLNIP-VP--FRAVPV-PTVSWHKDGK--E-VK-AS--D----------------------------R--LTM-KN-D-HISAHLEVPKSVHADAGVYTITLENKLGSA-TASINVK----
16470 --------------------PKVILRT-S-LEVKRGD----EIALD-AT--ISGSPY-PTITWIKDEN--V-IV-PE--EIKKRAAPPVRRKKGEAEEEEPFSLPLTER--LSI-NNSK-QGESQLRIRDSLRPDHGQYMIKVENDHGVA-KAPCSVS----
17184 ----------------PTIKLRLAVRG-DTIKVKAGE----PVNIP-AD--VTGLPM-PKIEWSKNEK--V-ID-KP--T--------------------D--TLNITK--EEV-S-RS-EAKTELSIPKAAREDKGTYTITASNRLGSV-FRNVHVE----
17589 --------PATDIQEVPEVFIDIGAQ--DCLVCKAGS----QVKIP-AV--IKGRPT-PKSSWEFDGK--A-KK-AM--K--------------------DGVHDIPED--AQL-E-TA-ENSSVIIIPECTRAHSGKYSITAKNKAGQK-TANCRVK----
17906 ---------------PPAIELKE-----F-MEVEEGT----DVNIV-AK--IKGVPF-PTLTWFKAPP--K-KP-DS--K---------------------EPVVYDTH--VNK-Q-VV-DDTCTLVIPQSRRSDTGLYSITAVNNLGTA-SKEMRLN----
18311 ----------------PDLQLDASVRD-R-IVVHAGG----VIRII-AY--VSGKPP-PTVTWSMNER--A-L---------------------------------PQE--AAI-E-TT-AISSSMVIKNCQRSHQGVYSLLAKNEGGER-KKTII------
18607 ----------------PTL--DLDFRD-K-LTVRVGE----SFALT-GR--YSGKPK-PKIDWFKDEA--D-VL-ED--D----------------------------R--THI-K-TT-PTTLALEKTKAKRSDSGKYCVVVENSTGSR-KGFCQVN----
19005 ----------------PELILDANMAR-E-QHIRVGD----TLRLS-AI--IKGVPF-PKVTWKKEDR--E-A---------------------------------PTK--AQI-D-VT-PVGSKLEIRNAAHEDGGIYSLTVENPAGTK-TVS--------
19297 ----------------PSVELDVKLI--EGLVVKAGT----TVRFP-AI--IRGVPV-PTAKWTTDGT--E-IK--T--D--------------------D-------H--YTV-E-TD-SFSSVLTIKNCLRKDTGEYQLTVSNAAGTK-TVAVHLT----
19695 ----------------PEVELDVTCRD-V-ITVRVGQ----TIRIL-AR--VKGRPE-PDITWSKEGK--V-LV-KD--K----------------------------R--VDL-I-HD-LPRVELQIKEAVRADHGKYIISAKNSSGHA-QGSAIVN----
19990 ----------------PVL--DLKLSG-V-LTVKAGD----TIRLE-AG--VRGKPF-PEVAWTKDKDATD-LT-RS--P----------------------------R--VKI-D-TS-AESSKFSLTKAKRSDGGKYVITATNPAGSF-VAYATVN----
20393 ----------------PEIDLDASMRK-L-VVVRAGC----PIRLF-AI--VRGRPA-PKVTWRKVGI--D-NV--------------------------------VRK--GQV-D-LV-DTMAFLVIPNSTRDDSGKYSLTLVNPAGEK-AVF--------
20688 ---------------PPKIL----MP--EQITIKAGK----KLRVE-AH--VYGKPN-PICKWKKGDD--E-VV--T--S--------------------S-------H--LAI-H-KA-DGSSVLIIKDVTRKDSGYYSLTAENSSGSD-TQKIKVT----
21082 ----------------PRIELSVEMK--SLLTVKAGT----NVCLD-AT--VFGKPM-PTVSWKKDTT--P-IK--Q--A--------------------E-------G--IKM-A-MK-RNLCTLELFSVNRKDSGDYTITAENSSGSK-SATIKLK----
22165 ----------------PTIVLDPTIKD-G-LTVKAGD----SIVLSAIS--ILGKPL-PKSSWSRAGK--D-IR-PS--D----------------------------I--AQI-T-ST-PTSSMLTVKYATRKDAGEYTITATNPFGTK-EEHVKVS----
23248 ----------------PRIMVDVRFK--DTITLKAGE----AFKLE-AD--VSGRPP-PTMEWTKDGK--E-LE--G--T--------------------G-------K--LEI-K-IA-DFSTHLINKDSSRTDSGAYILTATNPGGFA-KHIFNVK----
23647 ----------------PEIELDADLRK-V-VTIRACC----TLRLF-VP--IKGRPA-PEVKWAREHG--E-S---------------------------------LDK--ASI-E-ST-SSYTLLVVGNVNRFDSGKYILTVENSSGSK-SAFVNVR----
23937 ----------------PAFK---LLFN-T-FTVLAGE----DLKID-VP--FIGRPP-PAVTWHKDDI--P-LK-QT--T----------------------------R--VNA-ES-T-ENNSLLTIKEACREDVGHYTVKLTNSAGEA-TETLNVI----
24330 ----------------PRISMDPKYR--DTVVVQAGE----SFKID-AD--IYGKPI-PTTQWVKGDQ--E-LS--S--T--------------------A-------R--LEI-K-ST-DFATSLSVKDAVRVDSGNYILKAKNVAGEK-SVT--------
24729 ----------------PDIDLDLELRK-V-INIRAGG----SLRLF-VP--IKGRPT-PEVKWGKVDG--D-I---------------------------------RDA--AII-D-VT-SSFTSLVLDNVNRYDSGKYTLTLENSSGTK-SAFVT------
25019 ----------------PDVR---PAFS-S-YSVQVGQ----DLKIE-VP--ISGRPK-PSISWTKDGM--P-LK-QT--T----------------------------R--INV-TD-S-LDLTTLSIKETHKDDGGQYGITVSNVVGQK-TAAIEIIT---
25412 ----------------PRISMDPKFR--DTIVVNAGE----TFRLE-AD--VHGKPL-PTIEWLRGDK--E-IE--E--S--------------------A-------R--CEI-K-NT-DFKALLIVKDAIRIDGGQYILRASNVAGSK-SFPVNVK----
26101 ----------------PSFK---LPFN-T-YSVQAGE----DLKIE-IP--VIGRPR-PKISWVKDGE--P-LK-QT--T----------------------------R--VNV-EE-T-ATSTILHIKESSKDDFGKYSVTATNSAGTA-TENLS------
26494 ----------------PNASLDPKYR--DVIIVHAGE----TFVLE-AD--IRGKPI-PDIIWSKDGN--E-LE-ET--A--------------------A-------R--MEI-K-ST-LQKTTLIVKDCIRTDGGQYTLKLSNVGGTK-TIPIT------
26894 ----------------PEIELDADLRK-V-VTLRASA----TLRLF-VT--IKGRPE-PEVKWEKAEG--I-L---------------------------------TER--AQI-E-VT-SSYTMLVIDNVTRFDSGRYNLTLENNSGSK-TAFVNVR----
27168 PRQLGVPVIAKDIEIKPSVE---LPFN-T-FNVKAND----QLKID-IP--FKGRPQ-ATVAWKKDGQ--V-LR-ET--T----------------------------R--VNV-AS-S-KTVTTLSIKEASREDVGTYELCVSNTAGSI-TVPITVI----
27576 ----------------PRIMMDVKFR--DVIIVKAGE----VLKIN-AD--IAGRPL-PVISWAKDGV--E-IE--E--R--------------------A-------K--TEI-V-ST-DYTTTLTVKDCVRRDTGQYVLTLKNVAGTR-TMA--------
27963 ------PQVAKEREEEPVFDVDSEMRK-T-LIVKAGS----SFTMT-VP--FRGRPI-PNVSWSKPDT--D-L---------------------------------RTR--AYI-D-ST-DSRTLLTIENANRNDSGKYTLTIQNVLSAA-SMT--------
28659 ----------------PVIDLPLEYT--EVVKYRAGT----SVKLR-AG--ISGKPE-PTIEWYKDDK--E-LQ--T--N--------------------A-------L--VCV-E-NS-TDLASILIKDANRLNSGSYELKLRNAMGSA-SATIRVQ----
29058 ----------------PEIDLDVALRT-S-VIAKAGE----DVQLL-IP--FKGRPP-PTVTWRKDEK--N-LG-S------------------------------DTR--YSI-Q-NT-DSSSLLVIPQVTRNDTGKYILTIENGVGQP-KSSTVS-----
29350 ----------------PEVDLSEIPGA-Q-ISVRIGH----NVHLE-LP--YKGKPK-PSISWLKDGL--P-LK-ES--E----------------------------Y--VRF-SK-T-ENKITLSIKNSKKEHGGKYTVILDNAVCRN-SFPIT------
29745 ----------------PTVEFGPEYF--DGLVIKSGD----SLRIK-AL--VQGRPV-PRVTWFKDGV--E-IE--R--R--------------------M-------N--MEI-T-DV-LGSTSLFVRDATRDHRGVYTVEAKNVSGST-KAEVTVK----
30144 ----------------PELDIDANFKQ-T-HIVRAGV----SIRLF-IA--YQGRPT-PTAVWSKPDS--N-L---------------------------------SIR--ADI-H-TT-DSFSTLTVENCNRNDAGKYTLTVENNSGKK-SIT--------
30430 ------------PQIEPTADLTGITN--QLITCKAGS----TFTID-VP--ISGRPA-PKVTWKLEEM--R-LK--E--T--------------------D-------R--MSI-A-TT-KDRTTLTVKDSMRGDSGRYFLTLENTAGVK-TFTIT------
30833 ----------------PKAELDARLQG-DLVTIRAGS----DLVLD-AA--VGGKPE-PKIIWTKGDK--E-LD-L------------------------------CEK--ISL-Q-YT-GKRATAVIKYCDRSDSGKYTLTVKNASGTK-SVS--------
31233 ----------------PDLELADDLKK-T-VIVRAGA----SLRLM-VS--VSGRPS-PVITWSKKGI--D-L---------------------------------ANR--AII-D-NT-ESYSLLIVDKVNRYDAGKYTIEAENQSGKK-SATVLVK----
31525 ----------------PTIDLSTMPQK-T-IHVPAGR----PIELV-IP--ITGRPP-PTASWFFAGS--K-LR-E------------------------------SER--VTV-E-TH-TKVTKLTIRETTIRDTGEYTLELKNVTGTT-SETIKVI----
31923 ----------------PDYELDERYQE-G-VFVRQGG----VIRLT-IP--IKGKPF-PVCKWTKEGQ--D-I---------------------------------SKR--AMI-A-TS-ETHTELVIKEADRNDSGTYDLVLENKCGKK-TVYIKVK----
32322 ----------------PGVR---KEMA-D-VTTKLGE----AAQLS-CQ--IVGRPL-PDIKWYRFGK--E-LI-QS--R----------------------------K--YKM-SS-D-GRTHTLTVMTDEQEDEGVYTCVATNEVGEV-ESSSKLL----
32717 ----------------PHFK---EELR-N-LNVRYQS----NATLV-CK--VTGHPK-PIVKWYRQGK--E-II-AD--G--------------------L-------KYRIQ--EF-K-GGYHQLIIASVTDDDATVYQVRATNQGGSV-SGTASLE----
32817 ----------------PKT----LEGM-GAVHALRGE----VVSIK-IP--FSGKPD-PVITWQKGQD--L-ID-NN--G----------------------------H--YQV-IVTR-SFTSLVFSNGVERKDAGFYVVCAKNRFGID-QKTVELDVA--
33358 -----------------PVS---GQIM-H-AIGEEGG----YVKYV-CK--IENYDQSTQVTWYFGVR--Q-LE-NS--E----------------------------K--YEI-TY-E-DGVATMYVKDITKFDDGTYRCKVVNDYGED-SSYAELF----
33478 ----------------PEFT---LPLY-N-KTAYVGE----NVRFG-VT--ITVHPE-PRVTWYKSGQ--K-IK-PG--D--------------------D-------EKKYTF-ES-D-KGLYQLTINSVTTDDDAEYTVVARNKHGED-SCKAKLTVT--
33583 ----------------PMFK---RLLA-N-AECHEGQ----SVCFE-IR--VSGIPA-PTLKWEKDGQ--P-LS-LG--P----------------------------H--IEI-VH-EGLDYYALHIRDTLPEDTGYYRVTATNTAGST-SCQAHLQ----
34163 ----------------PRIT---LRMR-S-HRVPCGQ----NTRFI-LN--VQSKPT-AEVKWYHNGV--E-LQ-ES--S----------------------------K--IHY-TN-T-SGVLTLEILDCQTEDGGTYRAVCTNYKGEA-SDYATLDVT--
34350 ----------------ARIL---TKPR-S-ITVHEGE----SARFS-CD--TDGEPV-PTVTWLREGQ--V-VS-TS--A----------------------------R--HQV-TT-T-KYKSTFEISSVQASDEGNYSVVVENSDGKQ-EAQFTLT----
34507 ----------------PKIT---QSLK-A-EASKDIA----KLTCA-VE--SSALCA-KEVAWYKDGK--K-LK-EN--G----------------------------H--FQF-HYSA-DGTYELKIHNLSESDCGEYVCEVSGEGGTS-KTSFQFT----
34641 ----------------PVIV---TGLR-D-TTVSSDS----VAKFT-IK--VTGEPQ-PTITWTKDGK--A-IA-QG--S----------------------------KYKLSS---N--KEEFILEILKTETSDGGLYACTVTNSAGSV-SSSCKLT----
34923 ----------------PSFI---TQPR-S-QNINEGQ----NVLFS-CE--VSGEPS-PEIEWFKNNL--P-IS-IS--S----------------------------NISVSR---S--RNVYTLEIRNAAVSDSGKYTIKAKNFHGQC-SATASLT----
35118 ----------------PKIEALP---S-D-ISIDEGK----VLTVA-CA--FTGEPT-PEITWSCGGR--K-IQ-NQ--E-------------------------QQGR--FHI-E-NT-DDLTTLIIMDVQKQDGGLYTLSLGNEFGSD-SATVNIN----
```

| Class | PEVK | N2Bus | Cys Free Ig | C47 C63 Ig | C47 Ig | C63 Ig | BFG Ig | BF Ig | BG Ig | FG Ig |
|---|---|---|---|---|---|---|---|---|---|---|
| **Number of instances in N2BA (uniprot Q8WZ42-1)** | 1 | 1 | 48 | 2 | 3 | 4 | 18 | 16 | 6 | 7 |
| **Number of instances in N2B (uniprot Q8WZ42-3)** | 1 | 1 | 29 | 2 | 3 | 3 | 0 | 7 | 2 | 2 |
| **Modification** | None | SS | None | ST | ST | ST | SS | SS | SS | SS |

**Supplementary Table S1. Classification of human titin I-band regions used for Monte Carlo simulations.** N2Bus region and domains containing at least 2 out of the 3 cysteines of the triad (BFG, BG, BF and FG) are assumed to be able to form disulfides (SS). Domains with Cys47 and/or Cys63 are classified as substrates for S-thiolation (ST). As a simplification, domains that do not contain any of these structurally conserved cysteines are classified as Cys-free domains and are therefore not considered to be affected by redox modifications. Also, domains that could fall into both the disulfide bond and the S-thiolation categories were also classified as Cys Free. The PEVK region lacks cysteine residues and therefore is not the target of any oxidative modification.

| Element | Contour length (nm) | Kuhn length (nm) |
|---|---|---|
| PEVK (N2BA isoform) | 721 | 1.82 |
| PEVK (N2B isoform) | 68 | 1.82 |
| Reduced N2Bus | 230 | 1.32 |
| Oxidized N2Bus | 125 | 1.32 |
| Folded Ig | 4 | 20 |
| Unfolded Ig (reduced) | 35.2 | 1.32 |
| Unfolded Ig (BG disulfide) | 10.8 | 1.32 |
| Unfolded Ig (BF disulfide) | 15.2 | 1.32 |
| Unfolded Ig (FG disulfide) | 31.2 | 1.32 |

**Supplementary Table S2. Mechanical parameters of the different elements in the titin model**. These parameters were obtained from the literature (*19*, *21*, *23*) and used for calculating the length of the protein according to the FJC model. Regarding oxidized N2Bus, we considered the most frequently established disulfides *in vitro*, Cys100-Cys367 and Cys7-Cys264 (*23*). Domains with the BFG triad behave according to the particular disulfide formed at each simulation point. Folded Ig domains have the same contour length and Kuhn length regardless of their oxidative status.

| | Unfolding | | Folding | |
|---|---|---|---|---|
| **Element** | **Δx (nm)** | **$\alpha_0$ ($s^{-1}$)** | **Δx (nm)** | **$\alpha_0$ ($s^{-1}$)** |
| Reduced Ig domain containing no more than one BFG cysteine | 0.25 | 8e-5 | -2.2 | 1.2 |
| Reduced Ig domain containing at least two BFG cysteines | 0.12 | 1e-4 | -2.2 | 0.05 |
| S-thiol in C47 and C63 | 0.25 | 1e-2 | -2.2 | 1.2 |
| S-thiol in C47 | 0.25 | 8e-5 | -2.2 | 1.2 |
| S-thiol in C63 | 0.25 | 1e-2 | -2.2 | 1.2 |
| BG disulfide | 0.12 | 1e-3 | -2.2 | 1.4 |
| BF disulfide | 0.12 | 1e-3 | -2.2 | 1.4 |
| FG disulfide | 0.12 | 1e-3 | -2.2 | 0.05 |

**Supplementary Table S3. Transition rates for unfolding and refolding reactions.** Domains with the BFG triad behave according to the particular disulfide formed at each simulation point.

| **Transition** | **$\Delta x$ (nm)** | **$\alpha_0$ ($s^{-1}$)** |
|---|---|---|
| BG → BF | 0.023 | 0.012 |
| BG → FG | 0.023 | 0.015 |

**Supplementary Table S4. Transition rates for isomerization reactions**.

| Element | Reduced Ig | Ig containing a disulfide | S-thiol at C47 and C63 | S-thiol at C47 | S-thiol at C63 |
|---|---|---|---|---|---|
| **Folding fraction** | 0.6 | 0.6 | 0.054 | 0.33 | 0.37 |

**Supplementary Table S5. Folding fraction of titin domains.** Folding probabilities calculated using Bell's model are multiplied by the folding fractions to obtain the final folding probability.

### Supplementary references

75. C. Neagoe, C. A. Opitz, I. Makarenko, W. A. Linke, Gigantic variety: expression patterns of titin isoforms in striated muscles and consequences for myofibrillar passive stiffness. *J Muscle Res Cell Motil* **24**, 175-189 (2003).
76. S. B. Smith, L. Finzi, C. Bustamante, Direct mechanical measurements of the elasticity of single DNA molecules by using magnetic beads. *Science* **258**, 1122-1126 (1992).
77. G. I. Bell, Models for the specific adhesion of cells to cells. *Science* **200**, 618-627 (1978).
